# Determinants of haplotype phasing accuracy in long-read human genome sequencing

**DOI:** 10.64898/2026.05.04.722832

**Authors:** Nikhita Damaraju, F. Graeme Frost, Jiayu Fu, David A. D’Onofrio, Joy Goffena, Sophie HR Storz, Zachary B. Anderson, Trent Prall, Miranda PG Zalusky, May Christine V. Malicdan, David R. Adams, Danny E. Miller

## Abstract

Accurate haplotype phasing is critical for interpreting human genetic variation. Long-read whole-genome sequencing has emerged as a powerful approach for read-based phasing, particularly where parental DNA is absent, yet the determinants of phasing accuracy remain incompletely defined. Here, we evaluate haplotype phasing performance across sequencing technology, reference genome, read length, and coverage depth using Oxford Nanopore Technologies (ONT) and Pacific Biosciences (PacBio) data from two Genome in a Bottle reference samples (HG002 and HG005). In clinically relevant genes, alignment to the T2T-CHM13 (T2T) reference genome improves phasing performance relative to GRCh38, reducing mean gene–level phasing error rates by 3–9-fold. T2T alignment increases phase set NG50 and yields 1.5–2-fold more phased variant pairs. At similar read N50 values, ONT has a higher phasing error rate than PacBio in certain genes. Downsampling demonstrates that phasing error rates plateau at ∼20x coverage. Longer ONT read lengths reduce phasing error rates and extend phase set contiguity. Haplotype-resolved assemblies produce substantially higher phasing error rates than alignment-based phasing, demonstrating the advantage of an alignment-based approach. To enable per-variant-pair confidence assessment, we introduce PhaseQuality, a technology-specific stratification method that assigns confidence tiers to phased variants based solely on sequencing data. PhaseQuality accurately assigns 82–99% of known phasing errors to lower-confidence tiers, reducing error rates among high-confidence pairs to <0.5%. Together, these results demonstrate the primary technical determinants of long-read haplotype phasing accuracy and provide practical benchmarks for optimizing reference genome selection, coverage targets, and read length for long-read sequencing studies.

## INTRODUCTION

Haplotype phasing—the assignment of genetic variants to their parental chromosomes—is fundamental to the interpretation of human genetic variation (Tewhey et al. 2011). Accurate phase resolution enables identification of compound heterozygous variants, clarifies *cis*–*trans* relationships among pathogenic alleles, and informs allele-specific regulatory and therapeutic analyses (Browning and Browning 2011). Despite its importance, high-confidence haplotype resolution at genome scale remains challenging (Hofmeister et al. 2023).

Conventional phasing strategies fall into two broad categories: pedigree-based inference and population-based statistical approaches (Browning and Browning 2011; Snyder et al. 2015). Pedigree phasing requires parental data, which may be unavailable or costly to obtain in both research and clinical contexts. Population-based statistical approaches offer the most commonly used alternative, relying on linkage disequilibrium patterns derived from reference panels; however, these methods are often unreliable for rare or private variants and are subject to population representation biases (Browning and Browning 2011; Hofmeister et al. 2023; Browning et al. 2021). Read-based phasing offers a third avenue that requires neither parental data nor reference panels, reconstructing haplotypes directly from reads that span multiple heterozygous sites into contiguous blocks known as phase sets (Martin et al. 2016; Snyder et al. 2015). Short-read sequencing limits read-based phasing to closely spaced variants and has low resolution in repetitive and low-complexity regions (Garg et al. 2016; Del Gobbo and Boycott 2025; Li and Freudenberg 2014; Wang et al. 2020).

Long-read sequencing (LRS) technologies, including Oxford Nanopore Technologies (ONT) and Pacific Biosciences (PacBio), may overcome these limitations by generating reads spanning tens of kilobases, enabling read-based phasing across a far greater proportion of the genome without the need for parental samples (Martin et al. 2016; Shafin et al. 2021). While LRS is increasingly being adopted for clinical application (Miller et al. 2021, 2022; Nakamura et al. 2020; Samarasinghe et al. 2026; van der Lee et al. 2022; Li et al. 2025), phasing accuracy has not been systematically evaluated across factors that routinely influence bioinformatics pipelines such as sequencing depth, read length, platform, and genomic context (Kolmogorov et al. 2023; Zheng et al. 2022). Prior studies have assessed genome-wide phasing performance (Martin et al. 2016; Kolmogorov et al. 2023; Lin et al. 2022), but focused evaluation within clinically relevant genes is lacking. For example, while the use of T2T-CHM13v2.0 (T2T) assembly has been shown to improve alignment and variant calling in challenging regions compared to GRCh38 (Aganezov et al. 2022; Altemose et al. 2022; Nurk et al. 2022; Schmitz et al. 2025), its impact on phasing accuracy has not been characterized.

Here, we perform a comprehensive analysis of haplotype phasing accuracy in LRS data from two well-characterized Genome in a Bottle reference samples (HG002 and HG005). We compare ONT R10 Simplex and PacBio HiFi sequencing data aligned to GRCh38 and T2T, quantify the effects of sequencing depth and read length, and evaluate performance across genomic contexts and Online Mendelian Inheritance in Man (OMIM) genes. As our goal is to assess how accurately long reads preserve haplotype relationships between variants across genomic distances, we use pairwise phasing error rate as the primary accuracy metric (Snyder et al. 2015). This analysis defines the principal technical determinants of long-read phasing accuracy and establishes empirical benchmarks for optimizing haplotype resolution in genome-scale sequencing studies.

## RESULTS

### Phasing performance varies by reference genome, variant distance, and genomic context

We evaluated the pairwise phasing accuracy for single nucleotide variants (SNVs) using ONT and PacBio whole-genome sequencing data from HG002 and HG005 aligned to GRCh38 and T2T **(Table 1)**. Across both samples and platforms, alignment to T2T consistently improved phasing contiguity and accuracy relative to GRCh38. The phase set NG50, which indicates the continuous distance over which variants can be phased, was longer for ONT than for PacBio across both reference genomes, consistent with ONT having a wider read length distribution despite the two datasets having similar read N50 values **(Fig. S1)**. Alignment to T2T further increased phase set contiguity for both technologies, resulting in higher NG50 values and a 1.5–2-fold increase in phased heterozygous variant pairs relative to GRCh38 **(Table 1**; **Fig. 1A)**. These gains were observed across all genomic distance bins and were particularly pronounced in difficult-to-align regions. Phasing error rates increased with genomic distance between variant pairs and were consistently higher in GIAB-defined difficult-to-align regions—due to low complexity, homology, or technical challenges—compared to non-difficult regions **(Fig. 1B)**. However, T2T alignments reduced phasing error rates across both genomic contexts. In non-difficult regions, mean error rates were reduced to ≤0.37% across technologies with T2T, compared to 1.1–1.3% with GRCh38. Improvements were also observed in difficult regions, although error rates remained elevated relative to non-difficult regions.

**Figure 1.**
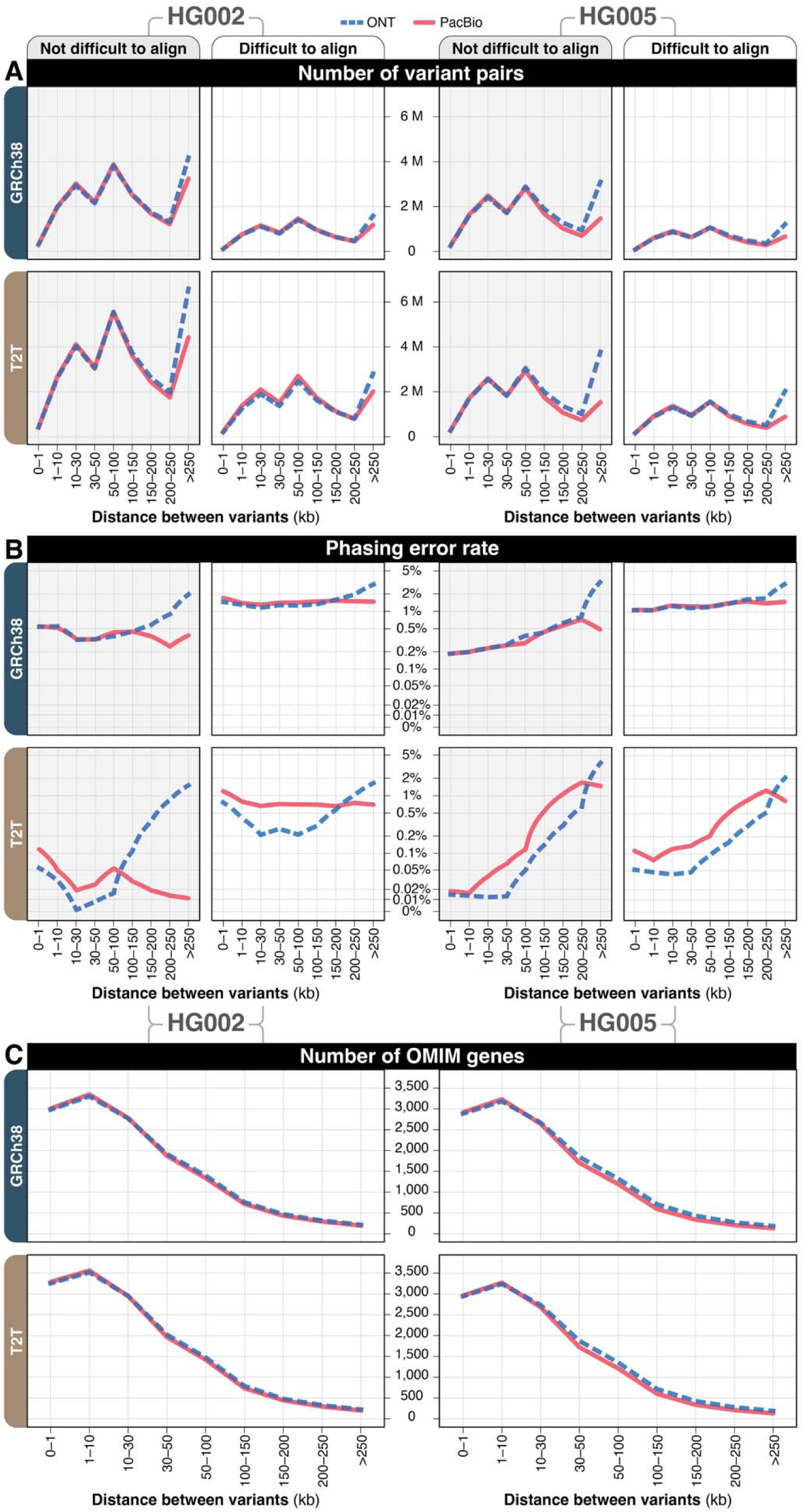
Number of variant pairs phased and phasing error rate as a function of variant distance, reference genome, and genomic context. **(A)** Number of high-quality phased variant pairs in HG002 and HG005 by distance between variants, stratified by sequencing technology, reference genome, and GIAB-defined genomic regions. **(B)** Phasing error rates in HG002 and HG005 by distance between variants, stratified by sequencing technology, reference genome, and GIAB-defined genomic regions. **(C)** Number of OMIM genes in HG002 and HG005 with heterozygous variant pairs phased by distance between variants for ONT and PacBio, stratified by reference genome.

**Table 1:** Summary sequencing and phasing metrics for ONT and PacBio aligned to GRCh38 and T2T reference genomes. Read N50 and depth of coverage represent input sequencing characteristics. Phase set NG50 indicates the continuous distance over which variants can be phased, where 50% of phased variants are in blocks of this size or larger.

| Sample | Platform | GRCh38 |  |  |  | T2T |  |  |  |
| --- | --- | --- | --- | --- | --- | --- | --- | --- | --- |
|  |  | Read N50 (kb) | Depth of coverage (x) | Phase setNG50 (kb) | Total number of heterozygous variant pairs | Read N50 (kb) | Depth of coverage (x) | Phase set NG50 (kb) | Total number of heterozygous variant pairs |
| HG002 | ONT | 22.3 | 41.2 | 595.7 | 28,791,614 | 22.6 | 40.7 | 622.2 | 44,061,444 |
|  | PacBio | 17.2 | 48.4 | 367.9 | 27,610,051 | 17.3 | 48.2 | 377.6 | 41,573,741 |
| HG005 | ONT | 20.9 | 37.6 | 428.4 | 22,080,971 | 21.1 | 37.2 | 443.6 | 26,547,181 |
|  | PacBio | 16.9 | 48.7 | 170.0 | 19,128,267 | 16.9 | 48.3 | 176.4 | 22,137,537 |

ONT produced more long-distance variant pairs than PacBio, particularly beyond 100–250 kb, reflecting its wider read length distribution **(Fig. 1A)**. This increase in long-range phasing contributed to modestly higher phasing error rates for ONT at the longest distance bins, indicating a tradeoff between phase set length and accuracy at extended genomic distances. The number of OMIM genes containing phased heterozygous variant pairs decreased with increasing variant distance for both platforms and reference genomes **(Fig. 1C)**. However, T2T alignments retained a greater number of OMIM genes with long-range phased variants, demonstrating improved haplotype resolution in clinically relevant loci.

To determine whether haplotype-resolved *de novo* assembly improves phasing accuracy relative to alignment-based methods, we compared diploid assemblies to read alignment-based phasing across both samples, technologies, and reference genomes. Assembly-based approaches substantially increased long-range haplotype contiguity. Beyond 200 kb, assemblies yielded 2–4-fold more phased variant pairs than alignment-based methods due to contig lengths exceeding individual read lengths **(Figs. S2A-B)**. However, this increase in contiguity was accompanied by markedly higher phasing error rates. Error inflation was most pronounced at extended variant distances, where assembly-based error rates frequently exceeded 30%, despite improved contiguity **(Figs. S2C-D)**. In contrast, alignment-based phasing maintained low error rates (<1% with T2T alignment) across most distance bins.

Together, these results demonstrate that reference genome choice, genomic context, and variant distance are the primary determinants of long-read phasing performance at gene scale. Additionally, the increased haplotype contiguity achieved through *de novo* assembly does not necessarily translate into improved phasing accuracy.

### Phasing Error Rates across OMIM Genes

To evaluate phasing performance at clinically relevant loci, we quantified gene-level phasing error rates across OMIM genes for each sequencing platform and reference genome **(Table 2)**. Alignment to T2T increased the number of OMIM genes containing phased heterozygous variant pairs and substantially reduced mean gene-level phasing error rates relative to GRCh38 in both samples and across both technologies. For HG002, 311 OMIM genes exhibited error rates >1% using ONT aligned to GRCh38 while 334 genes using PacBio HiFi had error rates >1%. Alignment to T2T reduced these counts to 56 (ONT) and 181 (PacBio). Similar trends were observed in HG005, where GRCh38 alignments yielded >1% error rates in 308–311 genes, compared to 30–45 genes when aligning to T2T **(Fig. 2)**. Across samples, T2T consistently reduced both the magnitude and prevalence of gene-level phasing errors.

**Figure 2.**
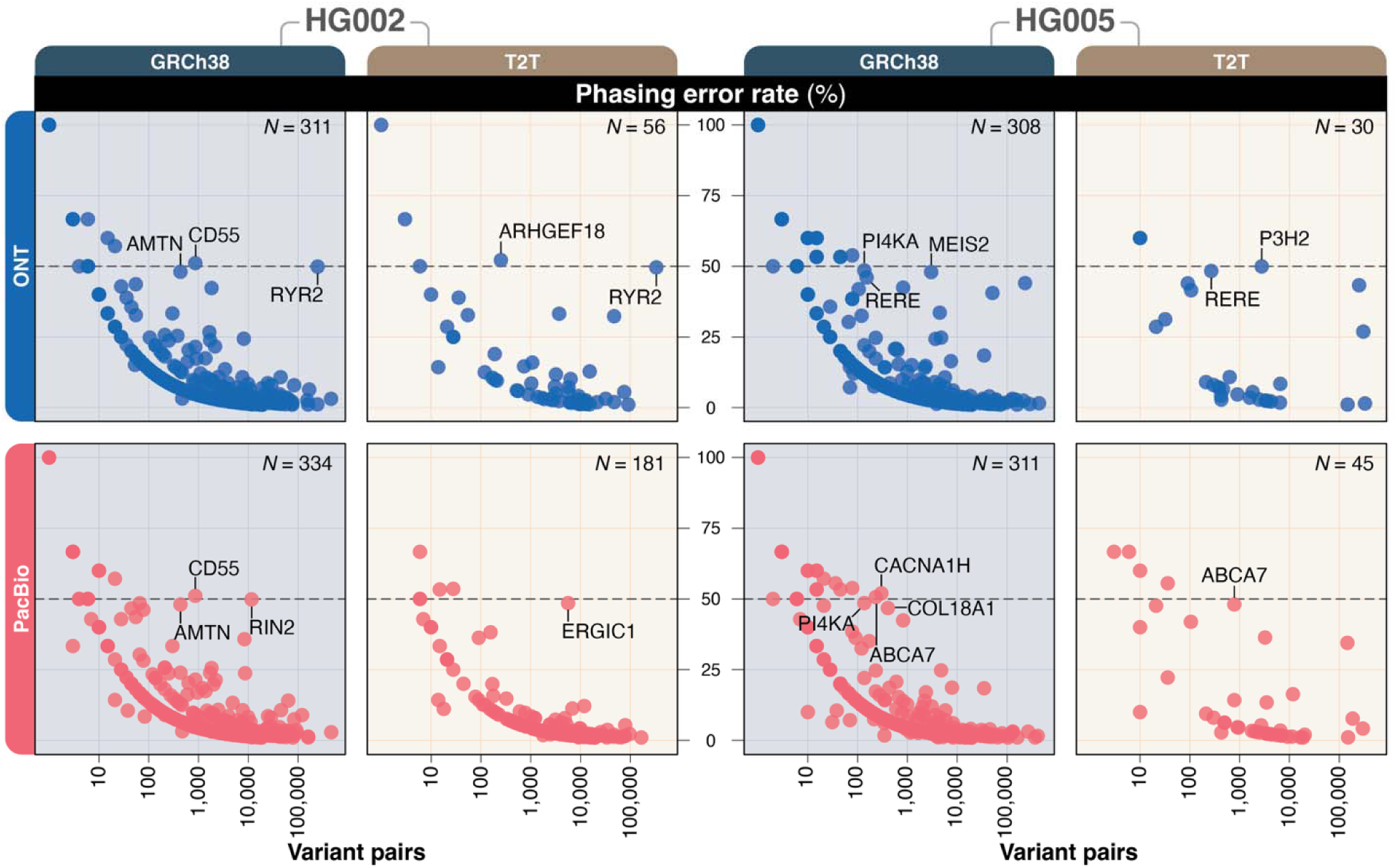
Gene-level phasing error rates in OMIM genes across sequencing technologies and reference genomes. Phasing error rate versus number of variant pairs for individual OMIM genes in HG002 (left) and HG005 (right) where each point represents a single gene with a phasing error rate greater than 1%. Genes with elevated phasing error rates and greater than 100 variant pairs are labeled, and the horizontal dashed line indicates 50% phasing error rate.

**Table 2:** Summary phasing metrics for OMIM genes by sequencing technology and reference genome. The number of OMIM genes with at least one heterozygous variant pair in a single phase set and mean phasing error rates are shown for ONT and PacBio HiFi aligned to the GRCh38 and T2T reference genomes in HG002 and HG005 samples.

| Sample | Platform | GRCh38 |  | T2T-CHM13v2.0 |  |
| --- | --- | --- | --- | --- | --- |
|  |  | Number of OMIM genes with at least 1 variant pair in a single phase set | Mean phasing error rate | Number of OMIM genes with at least 1 variant pair in a single phase set | Mean phasing error rate |
| HG002 | ONT | 3,536 | 1.14% | 3,733 | 0.23% |
|  | PacBio | 3,565 | 1.15% | 3,756 | 0.37% |
| HG005 | ONT | 3,466 | 1.34% | 3,503 | 0.15% |
|  | PacBio | 3,450 | 1.28% | 3,490 | 0.20% |

Eleven OMIM genes in HG002 and 13 genes in HG005 demonstrated phasing error rates exceeding 25% in at least one technology when aligned to GRCh38 (**Supplementary Table 1**). For the majority of these loci, T2T alignment substantially reduced error rates across both platforms. However, T2T alignment introduced elevated error rates at a small number of loci that were accurately phased on GRCh38. For example, error rates in *ARHGEF18* in HG002 increased with T2T alignment (52.2% ONT; 38.2% PacBio) compared to GRCh38 (0% ONT; 0% PacBio). This increase coincided with a larger phase set in T2T, resulting in a greater number of long-distance variant pairs incorporated into a single phase set **(Fig. 3A, S3)**. Phasing error rates in this locus rose sharply with inter-variant distance, reaching 100% beyond 50 kb, indicating that improved phase set contiguity can lead to long-range phasing inaccuracies that are not present with shorter phase sets (**Fig. S4**). This pattern was not observed in HG005 for this gene, where higher variant density supports more contiguous phase sets and enables accurate phasing across the locus, as reflected in 0% phasing error rates with T2T alignment **(Fig. 3A).**

**Figure 3.**
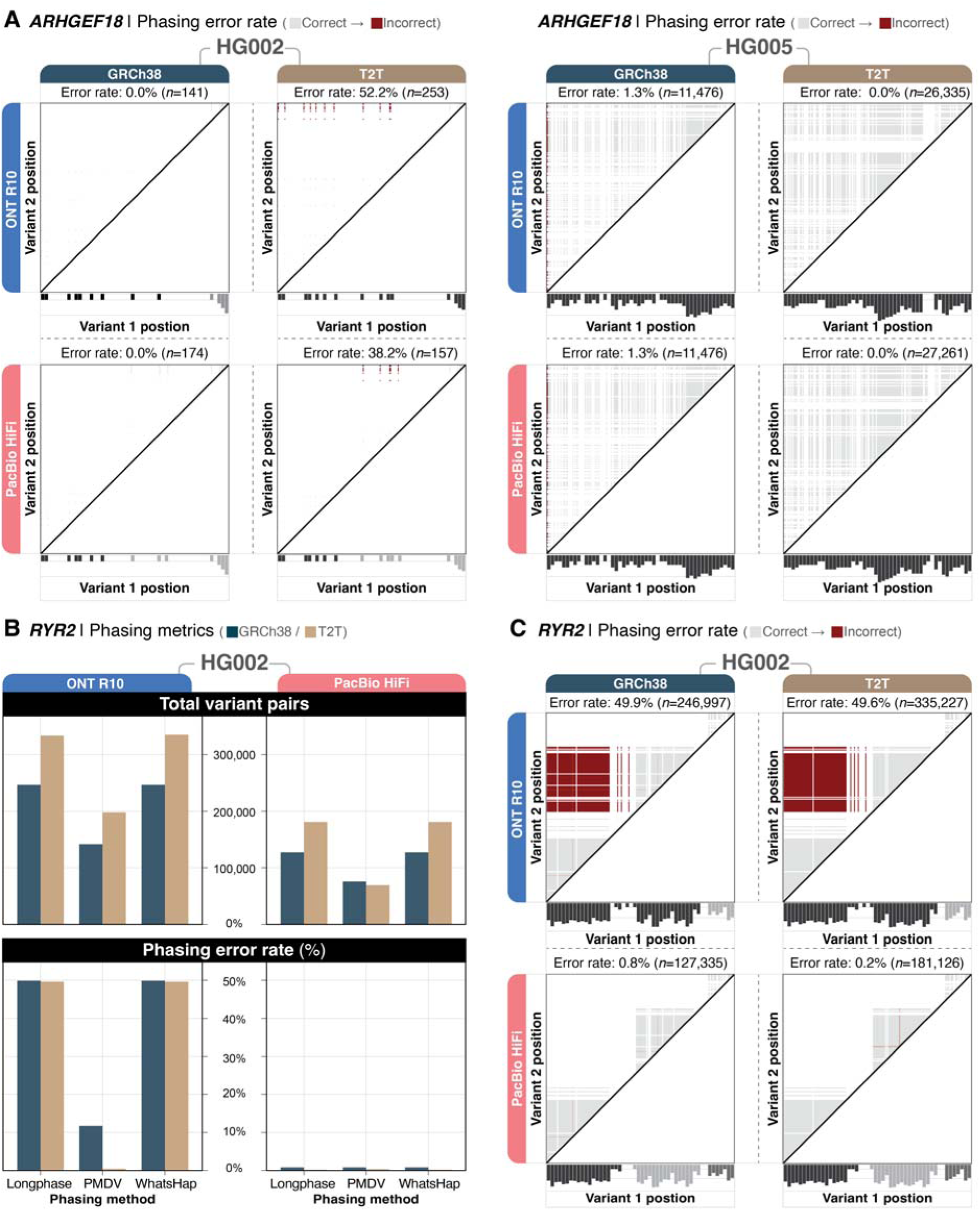
Phasing error rate analysis in *ARHGEF18* and *RYR2*. **(A)** Pairwise phasing error rate matrices for *ARHGEF18* in HG002 (left) and HG005 (right), showing variant-pair-level phasing accuracy across genomic positions for ONT R10 and PacBio HiFi aligned to GRCh38 and T2T; grey cells indicate correctly phased variant pairs and red cells indicate incorrectly phased variant pairs. Bottom graph represents SNV density and is colored by phase set. HG002 has very few variant pairs for this gene compared to HG005, demonstrating how increased SNV density reduces phasing error rate. **(B)** Comparison of phasing metrics for *RYR2* in HG002 across phasing methods; the top panel shows total variant pairs phased for each method and technology combination, and the bottom panel shows phasing error rates using Longphase, PMDV, and WhatsHap. **(C)** Pairwise phasing error rate matrices for *RYR2* in HG002, showing variant-pair-level phasing accuracy across genomic positions; grey cells indicate correctly phased variant pairs and red cells indicate incorrectly phased variant pairs. Bottom graph represents SNV density within the gene and is colored by phase set.

To assess the contribution of variant calling and phasing algorithms to gene-level variability, we compared three pipelines: 1) Clair3 + WhatsHap (primary), 2) Clair3 + Longphase, and 3) Pepper-Margin-DeepVariant (PMDV). We focused on *RYR2* as a representative example as it exhibited elevated phasing error rates in HG002 ONT data across both reference genomes while remaining consistently well-phased in PacBio HiFi. ONT aligned to GRCh38 exhibited high phasing error rates (∼49%) using Longphase and WhatsHap, while PMDV reduced error rates to 11.8%, albeit with fewer phased variant pairs due to shorter phase sets created by PMDV **(Fig. 3B)**. T2T alignment had no effect on Longphase or WhatsHap error rates (∼49.6%), whereas PMDV benefited substantially, with the error rate dropping to 0.5%. In contrast, PacBio HiFi exhibited consistently lower error rates (<1%) for *RYR2* across pipelines and reference genomes. This indicates technology-specific locus effects that result in shorter, but a greater number of phase sets. PacBio phasing segmented *RYR2* into three separate phase sets regardless of algorithm choice, eliminating the long-range variant pairs that drive error accumulation in the single extended ONT phase set **(Fig. 3C, S5)**.

*HYDIN* illustrated a different pattern: phasing error rates were ∼3.5% across all pipelines in GRCh38 for both technologies, whereas T2T alignment reduced error rates to near zero **(Fig. S6, S7)**. This suggests that reference genome representation, rather than algorithm selection, was the dominant determinant of phasing accuracy at this locus. Together, these analyses demonstrate that reference genome choice exerts a larger influence on gene-level phasing accuracy than variant calling or phasing algorithm selection in most OMIM genes. However, locus-specific interactions among reference representation, phase set contiguity, variant density, and sequencing technology can produce heterogeneous outcomes at individual genes.

Assembly-based phasing exhibited 3–6-fold higher mean gene-level error rates within OMIM genes compared to alignment-based approaches, despite improved contiguity (**Supplementary Table 2**). At high-error loci, T2T substantially reduced alignment-based error rates whereas assembly-based error rates improved only modestly, and alignment-based phasing outperformed assembly in 10–14% of OMIM genes with mean improvements of ∼30%, while assembly showed improved accuracy in ≤1.4% of genes (**Fig. S8–S11; Supplementary Table 3**). These findings indicate that increased haplotype contiguity achieved through de novo assembly does not translate into improved phasing accuracy at gene scale, with alignment-based approaches providing more reliable haplotype inference in clinically relevant genes, at least at the read N50 values assessed in this study (17-22kb N50).

### Effect of sequencing depth on phasing error rate

To quantify the impact of sequencing depth on phasing performance, ONT and PacBio HiFi datasets for HG002 and HG005 were downsampled to 10x, 20x, and 30x coverage and evaluated across variant distance bins and genomic contexts **(Table 3**; **Fig. 4)**. Phase set contiguity increased progressively with sequencing depth for both platforms and reference genomes. In both samples, phase set NG50 approximately doubled between 10x and 30x coverage, with longer phase sets consistently observed for ONT relative to PacBio and for T2T relative to GRCh38. However, the rate of improvement diminished above 20x coverage, indicating diminishing returns for additional sequencing depth.

**Figure 4.**
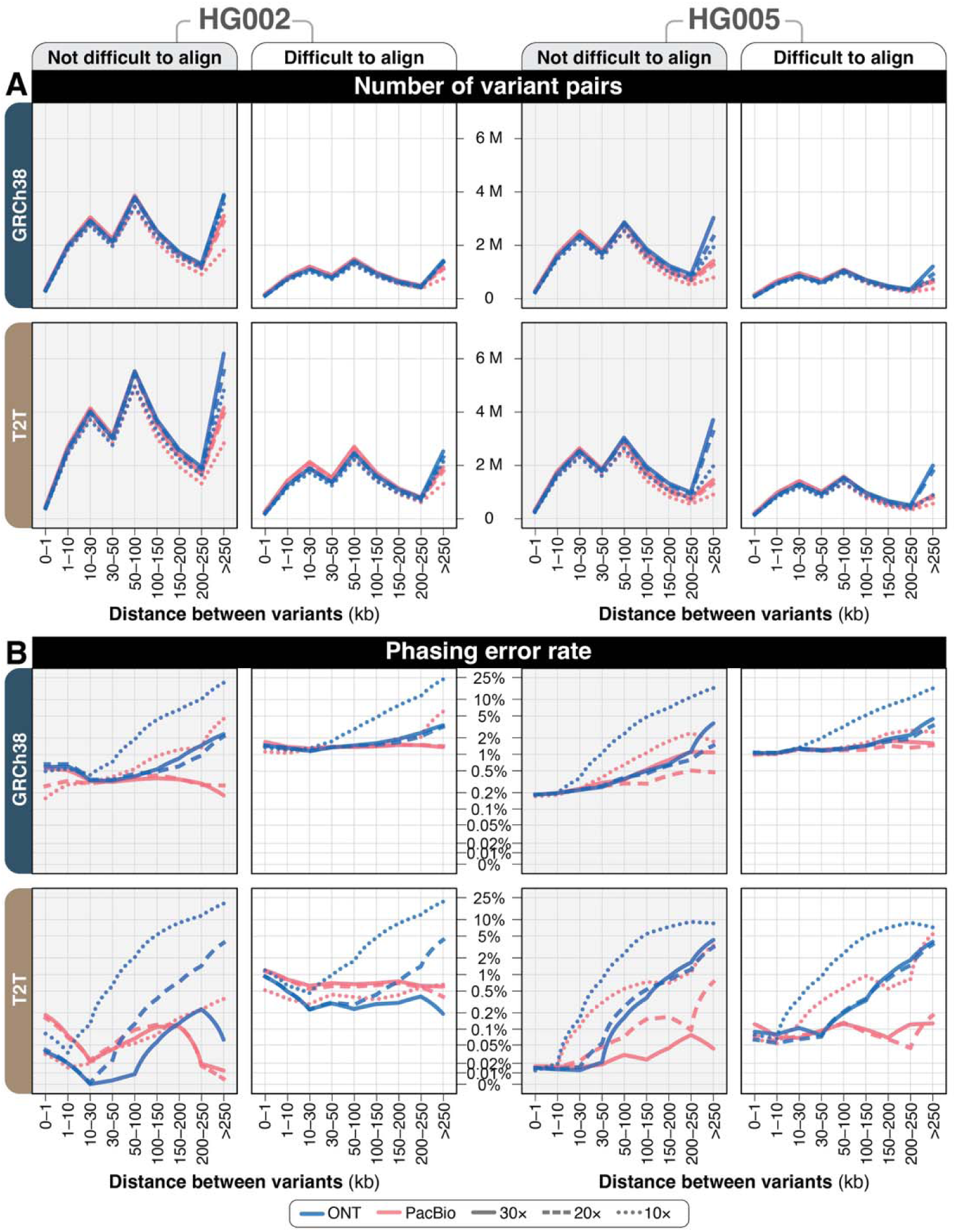
Effect of sequencing depth on number of variant pairs phased and phasing error rate. **(A,B)** Number of high-quality phased variant pairs in HG002 **(A)** and HG005 **(B)** by distance between variants for ONT and PacBio at varying coverage levels (10×, 20×, 30×), stratified by reference genome and GIAB-defined genomic regions. **(C,D)** Phasing error rates by distance between variants in HG002 **(C)** and HG005 **(D)** for ONT and PacBio at varying coverage levels (10×, 20×, 30×), stratified by reference genome and GIAB-defined genomic regions.

**Table 3:** Summary sequencing and phasing metrics for ONT and PacBio in HG002 and HG005 at varying coverage levels (10×, 20×, 30×) aligned to GRCh38 and T2T. Read N50 and depth of coverage represent sequencing characteristics. Phase set NG50 indicates the continuous distance over which variants can be phased and increases with sequencing coverage.

| Sample | Platform (coverage) | GRCh38 |  |  | T2T |  |  |
| --- | --- | --- | --- | --- | --- | --- | --- |
|  |  | Read N50 (kb) | Phase set NG50 (kb) | Total number of phased variant pairs | Read N50 (kb) | Phase set NG50 (kb) | Total number of phased variant pairs |
| HG002 | ONT (10x) | 22.4 | 364.9 | 15,780,790 | 22.5 | 378.5 | 37,941,412 |
|  | ONT (20x) | 22.4 | 470.6 | 17,037,630 | 22.5 | 493.6 | 41,846,846 |
|  | ONT (30x) | 22.4 | 540.3 | 17,372,997 | 22.5 | 565.7 | 43,225,050 |
|  | PacBio (10x) | 17.2 | 165.3 | 13,496,639 | 17.3 | 169.3 | 33,943,844 |
|  | PacBio (20x) | 17.2 | 270.9 | 15,996,616 | 17.3 | 275.6 | 39,413,314 |
|  | PacBio (30x) | 17.2 | 320.1 | 16,491,291 | 17.3 | 328.6 | 40,508,961 |
| HG005 | ONT (10x) | 20.9 | 273.6 | 25,802,319 | 21.1 | 282.0 | 20,648,987 |
|  | ONT (20x) | 20.9 | 338.2 | 27,802,288 | 21.1 | 349.1 | 25,066,479 |
|  | ONT (30x) | 20.9 | 391.3 | 28,252,349 | 21.1 | 408.3 | 26,212,903 |
|  | PacBio (10x) | 16.9 | 94.5 | 22,290,470 | 16.9 | 98.2 | 17,775,778 |
|  | PacBio (20x) | 16.9 | 137.2 | 26,094,548 | 16.9 | 142.1 | 20,875,452 |
|  | PacBio (30x) | 16.9 | 154.8 | 26,928,336 | 16.9 | 159.2 | 21,592,328 |

The number of high-quality phased variant pairs increased with coverage across all distance bins **(Fig. 4A–B)**. At 10x coverage, variant pair counts declined sharply beyond 100–250 kb, particularly for PacBio HiFi, consistent with reduced read support for long-distance phasing. In contrast, 20x and 30x datasets showed similar variant pair counts across most distance bins, with differences largely confined to the longest (>250 kb) intervals. Across all coverage levels, T2T alignments yielded 1.5–2-fold more phased variant pairs than GRCh38.

Phasing error rates were strongly influenced by coverage at long variant distances but plateaued at moderate coverage levels **(Fig. 4C–D)**. In GRCh38 alignments, 10x coverage exhibited elevated error rates beyond 30–50 kb, particularly for ONT, where long-distance error rates reached 20–30% in HG002. Increasing coverage to 20x substantially reduced error rates across distance bins, with minimal additional improvement at 30x. This plateau effect was observed in both non-difficult and difficult-to-align regions.

Alignment to T2T consistently reduced phasing error rates across all coverage levels. Notably, 10x coverage aligned to T2T achieved error rates comparable to or lower than 30x coverage aligned to GRCh38 in non-difficult regions. At 20x and 30x coverage, T2T alignments maintained low error rates across most variant distances, with remaining errors concentrated in difficult-to-align regions.

Together, these results demonstrate that ∼20x long-read coverage is sufficient to achieve near-optimal phasing accuracy in clinically relevant genes, while additional sequencing primarily increases phase set contiguity rather than substantially reducing error rates. Reference genome selection exerts a larger effect on phasing accuracy than increasing coverage beyond this threshold.

### Effect of read length on phasing performance

To evaluate the independent contribution of read length to phasing accuracy, we compared standard ONT datasets (read N50 ∼22 kb) to ONT datasets with substantially longer read lengths (N50 ∼85 kb), generated at comparable coverage depths **(Table 4)**. Increasing read length dramatically extended phase set contiguity. Phase set NG50 increased from ∼600 kb with standard ONT to 29–50 Mb with longer read length data, reflecting the ability of long reads to bridge extended haplotype intervals. Variant pair counts were similar across read-length conditions for inter-variant distances ≤250 kb but increased substantially for distances >250 kb with longer read length data **(Fig. 5A).**

**Figure 5.**
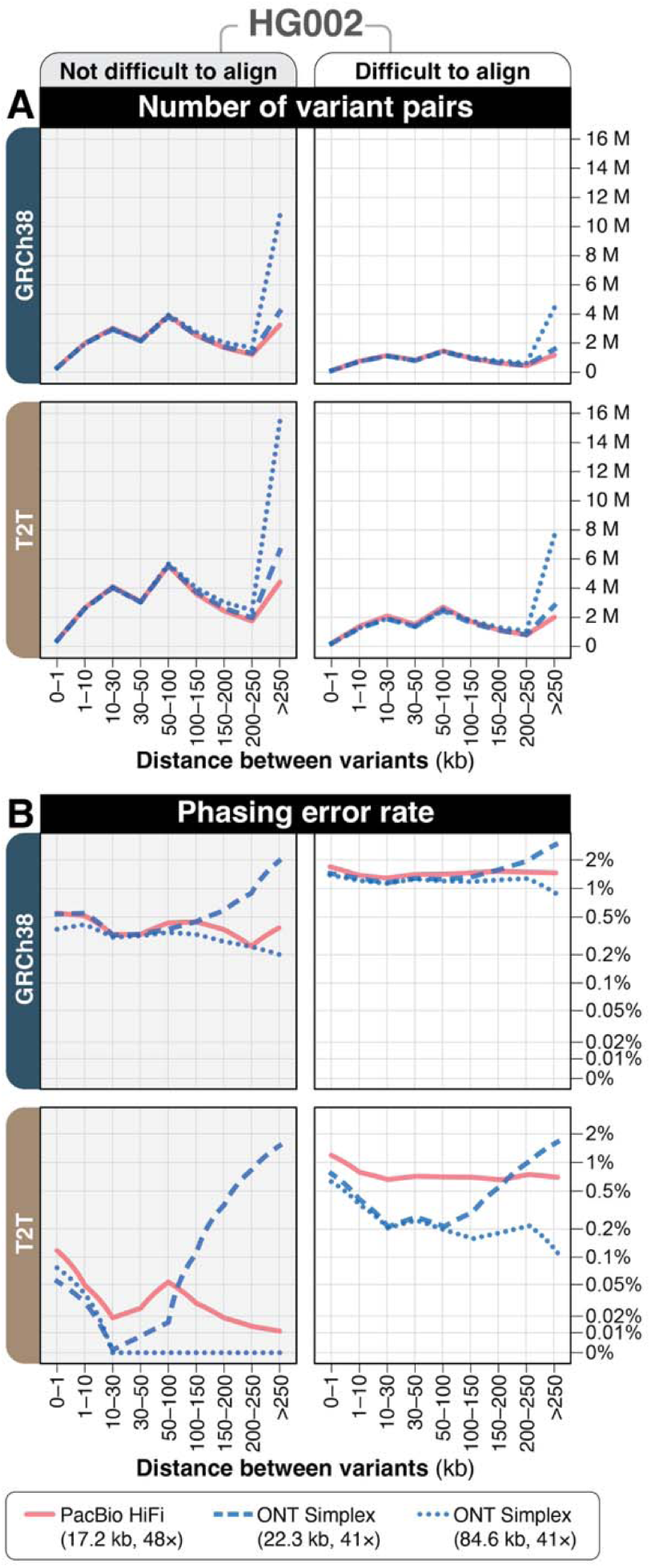
Effect of read length on number of variant pairs phased and phasing error rate for HG002. **(A,B)** Number of high-quality phased variant pairs **(A)** and phasing error rates **(B)** by distance between variants for ONT (read N50 ∼22 kb) and ONT (read N50 84.6–89.3 kb), stratified by reference genome and GIAB-defined genomic regions. PacBio data (read N50 17.2–17.3 kb) is shown for comparison.

**Table 4.** Summary sequencing and phasing metrics for ONT (read N50 ∼22 kb), ONT (read N50 ∼85 kb), and PacBio aligned to GRCh38 and T2T reference genomes. Read N50 and depth of coverage represent input sequencing characteristics. Phase set NG50 indicates the continuous distance over which variants can be phased, where 50% of phased variants are in blocks of this size or larger.

|  | GRCh38 |  |  | T2T-CHM13v2.0 |  |  |
| --- | --- | --- | --- | --- | --- | --- |
| Technology | Read N50 (kb) | Depth of coverage | Phase set NG50 | Read N50 (kb) | Depth of coverage | Phase set NG50 |
| ONT | 22.3 | 41.2 | 595.7 kb | 22.6 | 40.7 | 622.2 kb |
| ONT | 84.6 | 40.6 | 29.9 Mb | 89.3 | 41.2 | 49.7 Mb |
| PacBio | 17.2 | 48.4 | 367.9 kb | 17.3 | 48.2 | 377.6 kb |

Phasing error rates were modestly reduced with longer reads in both reference genomes **(Fig. 5B)**. In GRCh38 alignments, longer read N50 datasets showed lower error rates across most distance bins compared to standard ONT data. Alignment to T2T further reduced error rates, with longer read N50 data achieving ≤0.08% error in non-difficult regions and 0.11–0.65% in difficult-to-align regions.

Improvements in phasing accuracy with longer reads were most evident at extended genomic distances, where standard read lengths exhibited rising error rates. These findings indicate that read length and reference genome representation independently contribute to phasing accuracy, with longer reads primarily improving long-range haplotype contiguity while modestly reducing distance-dependent error accumulation.

### PhaseQuality: Technology-specific confidence stratification for long-read phasing

A key challenge in utilizing read-based phasing in clinical settings is being able to assess the accuracy without access to parental samples. To address this, we developed PhaseQuality, a stratification system that assigns confidence tiers (HighQ, MediumQ, LowQ) to each variant pair derived solely from a sample’s sequencing data.

We first trained separate random forest classifiers for each combination of long-read sequencing technology and reference genome on a balanced subset of HG002 that includes all incorrectly-phased variant pairs using six features derived from aligned files and reference genome annotation (**Methods**). Using feature importance analysis, we identified technology-specific error mechanisms. ONT models were driven by structural features: gap-to-read ratio, which is the maximum gap between variants divided by the longest aligned read in the phase set, and maximum gap between variants. On the other hand, PacBio models were dominated by difficult-to-align region annotations. **(Fig. S12)**. Based on these patterns, we developed technology-specific rules using ratio-based thresholds generalized across read lengths. ONT variant pairs were flagged as LowQ when gap-to-read ratio exceeded 0.5, or when the gap exceeds half the length of the longest aligned read in the phase set being evaluated. PacBio variant pairs were flagged as LowQ only when overlapping difficult-to-align regions and elevated gap-to-read ratio (>0.3) were present, preventing over-flagging in regions where high-accuracy reads maintain correct phasing.

To validate the performance, we applied PhaseQuality to the complete HG002 dataset and to HG005, demonstrating cross-sample generalization, with LowQ and MediumQ tiers capturing 82–99% (ONT) and 91–99% (PacBio) of phasing errors, reducing HighQ error rates to 0.005–0.35% (ONT) and 0.05–0.46% (PacBio), respectively **(Table 5).** These results indicate that phasing errors arise from distinct mechanisms—structural limitations in ONT data versus reference quality effects in PacBio. The utility of PhaseQuality is illustrated in two genes in HG005 (**Fig. 6**). In *MEIS2*, ONT R10 on GRCh38 exhibited a 48% error rate, with all 1,440 errors captured in the LowQ tier while 795 pairs (26%) remained in higher tiers with 0% error rate. Switching to T2T eliminated all ONT phasing errors, demonstrating how PhaseQuality identifies loci where reference choice resolves technology-specific failures (**Fig. 6A, B**). Conversely, *TACR3* exhibited PacBio-specific errors: ONT achieved 0% error on both references, while PacBio HiFi showed 12–13% error rates that persisted on T2T. PhaseQuality captured 86% of PacBio errors in the LowQ tier, reducing MediumQ error rates to ∼2% **(Fig. 6C,D**). Together, these examples demonstrate that PhaseQuality flags unreliable pairs across distinct error mechanisms unique to a technology and reference genome.

**Figure 6.**
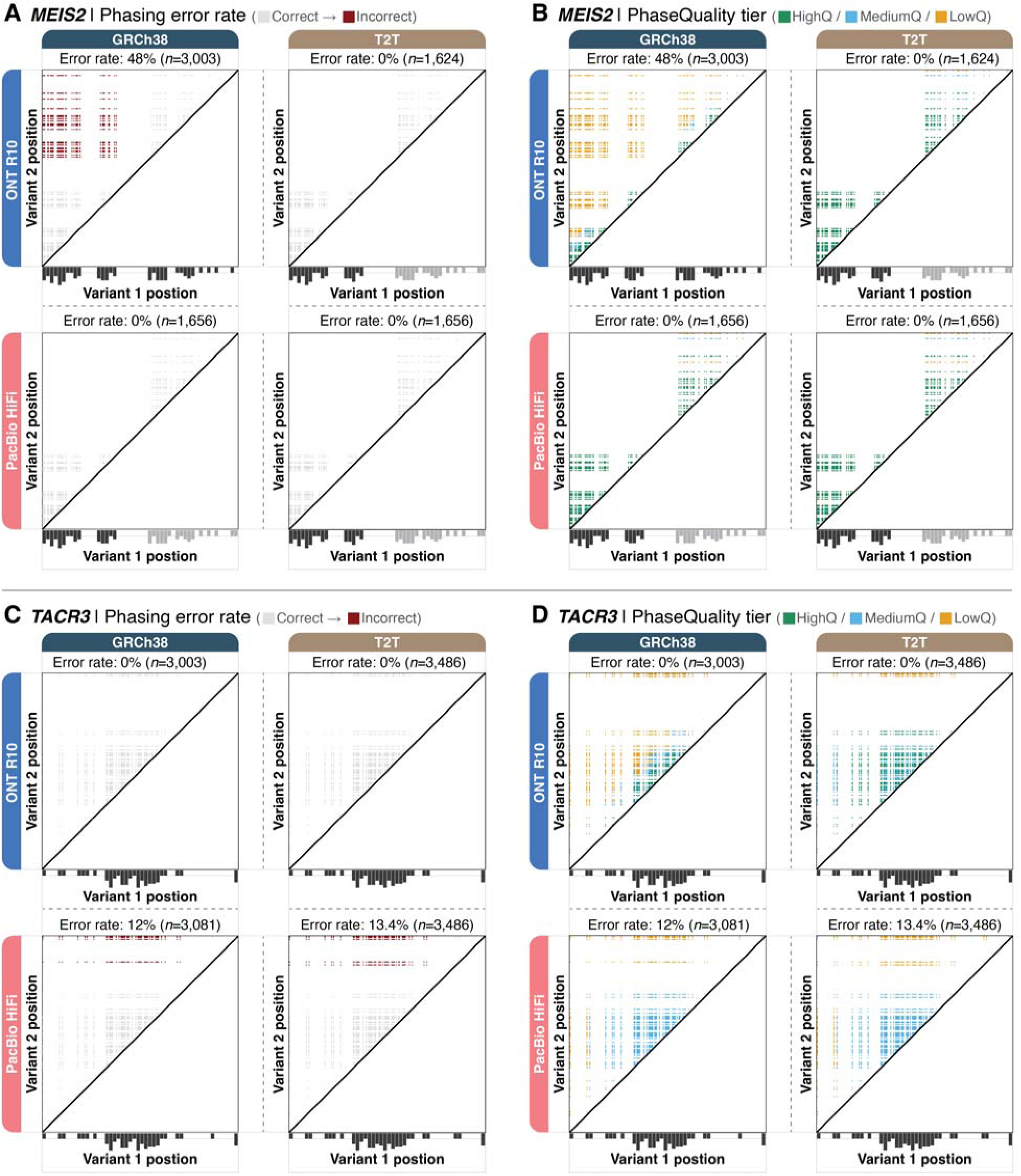
PhaseQuality stratification of *MEIS2* and *TACR3* variant pairs in HG005. Each cell represents a variant pair, with position along the gene shown on both axes. The diagonal line indicates self-comparisons; grey bars below each panel show variant density colored by phase set. Phasing error rate is denoted in parenthesis. **A and C.** Phasing error rates based on truth set (darker red indicates phasing errors). **B and D.** PhaseQuality tier assignments (HighQ: green, MediumQ: blue, LowQ: orange).

**Table 5.** PhaseQuality stratification of variant pairs across samples, technologies, and reference genomes. Variant pairs were assigned to LowQ, MediumQ, or HighQ tiers based on technology-specific rules using gap-to-read ratio and difficult-to-align region annotations. “% Error captured” indicates the percentage of all phasing errors within each technology-reference combination that fall into each tier. Rules were developed on a balanced training subset of HG002 and validated on the entirety of HG002 and HG005, demonstrating cross-sample generalization.

| Sample | Technology | Reference | PhaseQuality level |  |  |  |  |  |
| --- | --- | --- | --- | --- | --- | --- | --- | --- |
|  |  |  | LowQ |  | MediumQ |  | HighQ |  |
|  |  |  | % Error captured | Number of pairs | % Error captured | Number of pairs | % Error captured | Number of pairs |
| HG002 | ONT | GRCh38 | 52.1% | 3,913,800 | 29.6% | 9,640,151 | 18.3% | 15,237,663 |
|  |  | T2T | 87.3% | 6,618,383 | 12.2% | 15,889,051 | 0.5% | 21,554,010 |
|  | PacBio | GRCh38 | 13.6% | 1,443,756 | 77.6% | 22,595,800 | 8.8% | 3,570,495 |
|  |  | T2T | 18.4% | 2,806,509 | 79.1% | 33,822,864 | 2.4% | 4,944,368 |
| HG005 | ONT | GRCh38 | 91.8% | 16,957,013 | 6.2% | 2,701,583 | 2.1% | 2,422,375 |
|  |  | T2T | 24.2% | 2,612,669 | 74.5% | 11,389,034 | 1.3% | 12,545,478 |
|  | PacBio | GRCh38 | 24.2% | 1,467,900 | 70.5% | 15,133,884 | 5.3% | 2,526,483 |
|  |  | T2T | 37.4% | 2,081,020 | 61.1% | 17,417,097 | 1.4% | 2,639,420 |

## DISCUSSION

This study systematically defines the primary determinants of haplotype phasing accuracy in long-read whole-genome sequencing, with a focused evaluation of clinically relevant OMIM genes. Across two reference samples, two sequencing platforms, multiple coverage depths, and alternative phasing strategies, we find that reference genome representation exerts the largest influence on phasing accuracy, followed by sequencing depth, and read length. Alignment to T2T consistently reduced gene-level phasing error rates relative to GRCh38, with gains in phase set contiguity and the number of phased variant pairs that were particularly evident in difficult-to-align genomic regions. Notably, the impact of reference genome choice exceeded that of increasing coverage from 20x to 30x, demonstrating that improvements in reference representation directly translate into improved haplotype resolution at gene scale.

Although most OMIM genes showed reduced error rates with T2T alignment, a subset of loci exhibited persistent or context-dependent inaccuracies. In some genes, improved phase set contiguity exposed long-range phasing errors not detectable in shorter phase sets, illustrating that extended contiguity does not uniformly translate to improved accuracy. Locus-specific effects were observed across sequencing technologies and phasing pipelines, indicating that variant density, local homology, and genomic architecture modulate phasing performance, and that gene-level benchmarking is essential for interpreting haplotypes in clinically relevant loci.

Sequencing depth substantially influenced phasing performance at low coverage, but improvements plateaued above ∼20x, with minimal additional reduction in error rates at 30x. Additional coverage primarily extended phase set contiguity rather than improving per-pair accuracy, supporting ∼20x long-read coverage as a practical target for genome-scale haplotype resolution. Increasing ONT read length (N50 ∼85 kb) dramatically extended phase set NG50 into the megabase range and reduced distance-dependent error rates, though accuracy remained dependent on genomic context and reference representation. Together, these results indicate that read length and reference genome contribute independently to phasing performance.

Haplotype-resolved de novo assembly yielded substantially longer haplotype blocks and increased long-range variant pair counts relative to alignment-based methods. However, these gains were accompanied by markedly higher gene-level phasing error rates. In most OMIM genes, alignment-based phasing provided lower or comparable error rates, and assembly-based approaches rarely improved accuracy. These results indicate that extended contiguity achieved through assembly does not necessarily translate into improved haplotype accuracy in clinically relevant genes. Alignment-based phasing remains the more reliable strategy for routine gene-level haplotype inference under current long-read sequencing conditions, a finding consistent across both HG002 and HG005 despite their distinct ancestral backgrounds.

Our development of PhaseQuality provides a practical framework for identifying unreliable phased variant pairs without requiring parental validation data. By training on structural features of the sequencing data—gap-to-read ratio, maximum inter-variant gap, and difficult-to-align region annotations—PhaseQuality revealed that ONT and PacBio phasing errors arise from fundamentally distinct mechanisms: ONT errors are driven primarily by structural gaps between heterozygous sites that exceed the bridging capacity of individual reads, whereas PacBio errors concentrate in difficult-to-align regions where reference quality degrades alignment accuracy. This distinction has practical implications: for ONT data, PhaseQuality can flag long-range pairs likely to be unreliable even after reference upgrades, while for PacBio data, it identifies loci where switching to T2T would be expected to resolve most errors. In validation across both HG002 and HG005, the LowQ and MediumQ tiers captured 82–99% of errors, reducing HighQ error rates to below 0.5%. PhaseQuality thus complements reference genome selection and coverage optimization by providing per-pair confidence estimates that could inform downstream clinical interpretation, particularly for compound heterozygosity assessment where the accuracy of individual phase assignments directly affects diagnostic conclusions.

This study was restricted to autosomal single nucleotide variants and did not evaluate indels, structural variants, or sex chromosomes, which present additional phasing and variant detection challenges (Carey et al. 2022; Lin et al. 2022; Martin et al. 2016). While loci with elevated error rates were identified, the specific sequence features underlying these inaccuracies—such as segmental duplication content, local mappability, or structural complexity were not systematically modeled. PhaseQuality was trained and validated on two GIAB reference samples; its generalization to diverse ancestries, disease cohorts, and alternative library preparation methods remains to be established. Future work integrating sequence-level features may enable prediction of gene-specific phasing reliability and guide targeted algorithmic development. As sequencing technologies continue to evolve, improvements in resolution may alter platform-specific error profiles, potentially necessitating recalibration of the features used in PhaseQuality. Variant density plays a critical role in phasing success but may be systematically biased by reliance on a single linear reference genome; pangenome-based alignment and variant calling could provide a more representative characterization of variant density and its influence on phasing accuracy across diverse populations (Nyaga et al. 2025). Additionally, specialized loci such as multicopy gene families and structurally complex immune regions may require tailored phasing strategies (Chen et al. 2023; Mosbruger et al. 2025) beyond those evaluated here.

Together, these results define a hierarchy of determinants governing long-read haplotype phasing accuracy: reference genome representation exerts the largest effect, followed by sequencing depth and read length, while assembly-based strategies do not improve gene-scale accuracy under current implementations. PhaseQuality provides a complementary approach for flagging unreliable variant pairs based on technology-specific error mechanisms. These findings establish practical benchmarks for optimizing long-read sequencing studies and underscore the importance of accurate reference genomes and per-pair confidence assessment for reliable haplotype inference in genome-scale analyses.

## METHODS

### Data generation

PacBio HiFi sequencing data for HG002 generated on the Revio platform were obtained from the Genome in a Bottle Consortium (https://ftp.ncbi.nlm.nih.gov/ReferenceSamples/giab/data/AshkenazimTrio/HG002_NA24385_son/PacBio_HiFi-Revio_20231031/). PacBio HiFi data for HG005 generated using circular consensus sequencing (CCS) chemistry were downloaded from GIAB (https://ftp.ncbi.nlm.nih.gov/ReferenceSamples/giab/data/ChineseTrio/HG005_NA24631_son/PacBio_CCS_15kb_20kb_chemistry2/uBAMs/) as unmapped BAM files and merged into a single BAM using samtools merge. For ONT R10 Simplex sequencing, HG002 and HG005 cell lines were obtained from the Coriell Institute. Genomic DNA was extracted using the Puregene Cell Kit (Qiagen). DNA libraries were prepared using SQK-LSK114 (ONT) following the manufacturer’s instructions with the following exceptions: 1) End repair and A-tailing reaction time was extended to 30 minutes; 2) 3 µL Thermolabile Proteinase-K (New England Biolabs #P8111) in 12 µL of Q5 buffer (New England Biolabs #B9027) was added to the reaction mix after A-tailing and before the first bead cleanup to remove excess protein carryover from extraction; 3) Adapter ligation proceeded for one hour. Sequencing was conducted on a PromethION24 instrument using FLOPRO-114M flow cells with R10.4.1 pore chemistry. Basecalling was performed using Dorado 1.1.1 (ONT) using the SUP 5mCG_5hmCG model. Sequencing proceeded until pores were exhausted or 72 hours elapsed before flow cells were washed using the Flow Cell Wash Kit (EXP-WSH004) and reloaded with additional aliquot. ONT R10 Simplex data for HG002 with a N50 of 85 kb was obtained from ONT Benchmark Datasets (s3://ont-open-data/gm24385_2023.12/) and was included for comparison of read lengths. All datasets, whether downloaded or generated in-house, were converted to FASTQ format using samtools fastq to ensure consistency across all downstream bioinformatic analyses.

### Stratified BED files

BED files defining all stratified regions in GRCh38 (https://ftp-trace.ncbi.nlm.nih.gov/ReferenceSamples/giab/release/genome-stratifications/v3.3/GRCh38@all/GRCh38-all-stratifications.tsv) and CHM13v2.0 (https://ftp-trace.ncbi.nlm.nih.gov/ReferenceSamples/giab/release/genome-stratifications/v3.3/CHM13@all/CHM13-all-stratifications.tsv) were downloaded from the Genome in a Bottle (GiAB) Consortium. Stratified regions are broadly defined as regions of the genome that are difficult to align and/or call variants in, for reasons of low complexity, homology, or technical difficulty. More information can be found on the GiAB GitHub (https://github.com/genome-in-a-bottle/genome-stratifications).

### OMIM gene annotations

OMIM gene annotations for GRCh38 were obtained from https://www.omim.org/downloads/, filtered to retain only autosomal genes, and converted to BED format using a custom R script. For T2T, gene annotations were downloaded from UCSC https://hgdownload.soe.ucsc.edu/goldenPath/hg38/bigZips/genes/. Since OMIM does not provide CHM13-specific coordinates, gene boundaries were transferred from GRCh38 through coordinate-based matching, with manual curation for missing genes. Both annotation files were filtered to include only OMIM genes and converted to BED format using custom R scripts.

### HG002 and HG005 truth datasets

Reference phasing datasets were constructed using pedigree-based approaches specific to each reference genome. For HG002 and HG005 datasets aligned to GRCh38, high-confidence small variant draft benchmark VCF files for the proband, mother, and father were obtained from the Genome in a Bottle Consortium (https://ftp.ncbi.nlm.nih.gov/ReferenceSamples/giab/data) and filtered to retain only single nucleotide variants. Parental origin was assigned to each heterozygous variant in the proband using a custom R script that applied Mendelian inheritance principles, with exclusion of *de novo* mutations and variants that could not be unambiguously phased.

For HG002 aligned to T2T, the VCF from the Telomere-to-Telomere Consortium’s Q100 project (https://github.com/marbl/HG002) created using diploid assemblies was used as the truth set. For HG005 aligned to T2T, pedigree phasing was performed independently for ONT R10 Simplex and PacBio HiFi to determine parent-of-origin for all heterozygous variants. Parental samples (HG006 and HG007) for ONT R10 Simplex were sequenced and processed using identical protocols as HG005. Parental PacBio HiFi data were obtained as unmapped BAM files from GIAB (https://ftp.ncbi.nlm.nih.gov/ReferenceSamples/giab/data/ChineseTrio/HG005_NA24631_son/PacBio_CCS_15kb_20kb_chemistry2/uBAMs/) and processed through the same alignment and variant calling pipeline as HG005. A high-confidence phasing truth set was constructed by retaining only variants with concordant parental assignments across both platforms.

### Alignment to reference genome

Minimap2 v2.28(Li 2018) was used to align FASTQ files to the reference genome using the --map-ont flag for ONT datasets and --map-hifi for PacBio datasets. Samtools v1.21 (Danecek et al. 2021) was used to convert the aligned files from SAM format to BAM, sort by genomic coordinates, and index.

### Variant calling

Small variant calling was performed using Clair3 1.0.8 (Zheng et al. 2022) with technology-specific models. For ONT data, the r1041_e82_400bps_sup_v400 model was used with the --platform=ont parameter. For PacBio data, the hifi_revio model was applied with the --platform=hifi parameter. Variant calling was executed independently for each BAM file aligned to GRCh38 and T2T reference genomes. Clair3 1.0.8 was run with the --use_whatshap_for_final_output_phasing flag to generate phased VCF outputs.

### Polishing phase calls

Following variant calling, phasing was refined using WhatsHap (Martin et al. 2016) to improve phase set contiguity and accuracy. VCF files were first filtered to retain only single nucleotide variants (SNVs) and split by chromosome. WhatsHap phase was then executed independently for each chromosome to speed up computation using the --ignore-read-groups flag, with the original aligned BAM file and reference genome as inputs. Chromosome-specific phased VCFs were merged and sorted using bcftools sort (Danecek et al. 2021), then compressed and indexed with tabix to generate the final polished phased VCF.

### Variant filtering and annotation

Phased VCF files were filtered to retain high-quality heterozygous variants meeting the following criteria: phasing quality (QUAL) ≥15, phased heterozygous genotype (0|1 or 1|0) and autosomal location (chr1-chr22). Filtered variants were intersected with OMIM gene annotations using bedtools v2.31 intersect (-wa -wb flags) to assign gene names. Variants were further annotated by intersection with GIAB-defined difficult-to-align regions, with a binary flag (YES/NO) indicating genomic context. Final annotated VCF files included OMIM gene assignments and difficult region status for systematic comparison of phasing performance across samples, technologies, and reference genomes.

### Pairwise phasing error rate calculation

The pairwise phasing accuracy was evaluated by comparing predicted phase relationships (cis or trans) between heterozygous variant pairs across genomic distances to ground truth data. Variant pairs were stratified by genomic distance (nine bins: 0-1 kb, 1-10 kb, 10-30 kb, 30-50 kb, 50-100 kb, 100-150 kb, 150-200 kb, 200-250 kb and >250 kb) and genomic context (GIAB difficult-to-align regions). Variant pairs were annotated as difficult-to-align if one or both variants in the pair overlapped a GIAB difficult-to-align region. For gene-level analysis, variant pairs were grouped by OMIM gene and error rates were calculated per gene. Only variant pairs where both variants were in the truth set and phased in the same phase set were evaluated. All analyses were performed using custom R 4.2.1 scripts with data.table, dplyr, and readr packages. Phasing error rate was calculated as the proportion of incorrectly phased pairs within each stratum:

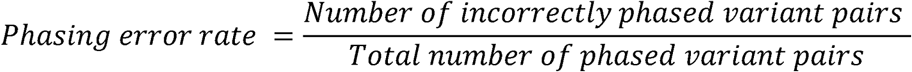

### Haplotagging for visualizing in IGV

To enable haplotype-aware visualization of phased variants in the Integrative Genomics Viewer (IGV), BAM files were haplotagged using longphase v1.7.3 (Lin et al. 2022). The haplotag module assigned haplotype information (HP tag) to individual reads based on their support for phased variants in the polished VCF files. Haplotagging was performed using the longphase haplotag command with a quality threshold of 1 and 20 threads. Output BAM files containing haplotype tags were indexed using samtools v1.21 to enable efficient visualization.

### Baseline metrics

Read N50 and depth of coverage were calculated using cramino 0.14.1 (De Coster & Rademakers, 2023). Phase set NG50 was calculated using whatshap stats.

### Coverage-based downsampling analysis

To generate downsampled datasets for coverage analysis, samtools v1.21 was used with a fixed random seed (2048) to ensure reproducibility. Reads were randomly selected using the --subsample option. For each sample, the subsampling fraction was calculated based on the coverage of the original BAM file calculated using cramino, such that the resulting datasets achieved target depths of 10×, 20×, and 30× coverage which were verified using cramino.

### Alternate variant callers and phasing algorithms

To compare phasing calling with alternate phasing software, longphase (v1.7.3) was used to phase variant call files generated by Clair3. Longphase performs haplotype phasing by identifying informative heterozygous SNPs and extending phase set through read-backed phasing of long-read sequencing data. Phasing was performed using the --ont flag for ONT data and the --pb flag for PacBio data. PEPPER-Margin-DeepVariant (PMDV) r0.8 (Shafin et al. 2021), an integrated variant caller and phasing software, was also used for comparison. PMDV combines PEPPER for candidate variant detection, Margin for read realignment, and DeepVariant’s deep learning model for variant calling, with built-in phasing capabilities. PMDV was run using Docker with the --ont_r10_q20 flag for ONT data and the --hifi flag for PacBio data.

Both tools were run on aligned BAM files for both ONT and PacBio datasets against GRCh38 and T2T reference genomes. Following phasing with both Longphase and PMDV, the resulting phased VCF files were used with longphase haplotag to assign haplotype information to individual reads in the aligned BAM files, enabling visualization in IGV.

### Haplotype-resolved de novo assembly

Haplotype-resolved assemblies were generated for HG002 and HG005 using technology-specific assemblers. For PacBio data, hifiasm v0.25.0-r726 (Cheng et al. 2021) was used with default parameters optimized for HiFi reads. For Oxford Nanopore R10 Simplex data, hifiasm-ont (Cheng et al. 2026) was used to generate phased assemblies. Both assembly workflows produced haplotype-specific contigs (.hap1.p_ctg.gfa and .hap2.p_ctg.gfa), which were converted to FASTA format and aligned to reference genomes (GRCh38 and T2T-CHM13v2.0) using minimap2.28. Dipcall (v.0.3) (Li et al. 2018) was used to perform variant calling for each haplotype-resolved assembly against GRCh38 and T2T-CHM13v2.0 reference genomes. Dipcall output VCF files contained phased variants with haplotype-specific genotypes. To ensure reliable phasing, variant pairs were generated only when both variants shared identical contig coverage patterns on haplotype 1 and haplotype 2, ensuring that phasing relationships were derived from continuous assembled sequences rather than potentially misaligned or chimeric regions. OMIM gene annotations and GIAB difficult-to-align region classifications were applied following the same workflow as alignment-based phasing, enabling direct comparison between assembly-based and alignment-based phasing approaches.

### PhaseQuality score development

To stratify variant pairs by phasing confidence, we developed PhaseQuality, a rule-based classification system assigning each pair to HighQ, MediumQ, or LowQ tiers. We computed six features for each variant pair: gap-to-read ratio (maximum gap between variants divided by the longest aligned read in the phase set), block-to-read ratio (phase set size divided by the longest aligned read), maximum gap between variants, variant density between variants, phase set size truncated to gene boundaries, and whether either variant overlapped a difficult-to-align region (segmental duplications or low mappability regions). Features were derived from haplotagged BAM files, phased VCFs, and Whatshap phase set annotations. To identify predictive features and inform threshold selection, we trained random forest classifiers using scikit-learn v1.3.0 (Pedregosa. 2011) with 100 trees and balanced class weights on a balanced subset of HG002 containing a subset of correct pairs and all incorrect pairs in a 2:1 ratio. Feature importance rankings were used to derive technology-specific rules using ratio-based thresholds that generalize across read lengths. For ONT, pairs were assigned LowQ when gap-to-read ratio exceeded 0.5 or when overlapping difficult regions with ratio exceeding 0.3; MediumQ when gap-to-read ratio exceeded 0.25 or when overlapping difficult regions; and HighQ otherwise. For PacBio, pairs were assigned LowQ when both overlapping difficult regions and having gap-to-read ratio exceeding 0.3, or when gap-to-read ratio exceeded 1.0; MediumQ when overlapping difficult regions, gap-to-read ratio exceeded 0.5, or block-to-read ratio exceeded 5; and HighQ otherwise. Rules were validated on the complete HG002 and HG005 datasets.

### Code availability

The code associated with this work is open source and can be found at https://github.com/millerlaboratory/lrs_phasing_accuracy_evaluation

## ACKNOWLEDGEMENTS

DEM is supported by NIH grant DP5OD033357. We would like to thank Angela Miller for expert assistance with editing and figure preparation. This study was supported by the Intramural Research Programs of the National Institutes of Health, including National Human Genome Research Institute. The contributions of the NIH authors are considered Works of the United States Government. The findings and conclusions presented in this paper are those of the authors and do not necessarily reflect the views of the NIH or the U.S. Department of Health and Human Services.

## AUTHOR CONTRIBUTIONS

Conceptualization: N.D., D.E.M. Data curation: N.D., J.G., S.H.R.S., Z.B.A., T.P., and M.P.G.Z. Formal analysis: N.D., F.G.F., J.F., and D.D. Investigation: J.G., S.H.R.S., Z.B.A., T.P., and M.P.G.Z. Methodology: N.D., F.G.F., and D.E.M. Resources: D.E.M. Supervision: D.E.M. Visualization: N.D. Writing—original draft: N.D., F.G.F., and D.E.M. Writing—review and editing: N.D., F.G.F., J.F., D.D., J.G., S.H.R.S., Z.B.A., T.P., M.P.G.Z., M.C.V.M., D.R.A., and D.E.M.

## REFERENCES

Aganezov S, Yan SM, Soto DC, Kirsche M, Zarate S, Avdeyev P, Taylor DJ, Shafin K, Shumate A, Xiao C, et al. 2022. A complete reference genome improves analysis of human genetic variation. Science 376: eabl3533.

Altemose N, Logsdon GA, Bzikadze AV, Sidhwani P, Langley SA, Caldas GV, Hoyt SJ, Uralsky L, Ryabov FD, Shew CJ, et al. 2022. Complete genomic and epigenetic maps of human centromeres. Science 376: eabl4178. https://www.science.org/doi/10.1126/science.abl4178 (Accessed February 10, 2026).

Browning BL, Tian X, Zhou Y, Browning SR. 2021. Fast two-stage phasing of large-scale sequence data. The American Journal of Human Genetics 108: 1880–1890. https://linkinghub.elsevier.com/retrieve/pii/S0002929721003049 (Accessed August 3, 2023).

Browning SR, Browning BL. 2011. Haplotype phasing: Existing methods and new developments. Nat Rev Genet 12: 703–714. https://www.ncbi.nlm.nih.gov/pmc/articles/PMC3217888/ (Accessed April 6, 2023).

Carey SB, Lovell JT, Jenkins J, Leebens-Mack J, Schmutz J, Wilson MA, Harkess A. 2022. Representing sex chromosomes in genome assemblies. Cell Genomics 2: 100132. https://www.sciencedirect.com/science/article/pii/S2666979X22000611 (Accessed March 17, 2026).

Chen X, Harting J, Farrow E, Thiffault I, Kasperaviciute D, Hoischen A, Gilissen C, Pastinen T, Eberle MA. 2023. Comprehensive SMN1 and SMN2 profiling for spinal muscular atrophy analysis using long-read PacBio HiFi sequencing. Am J Hum Genet 110: 240–250. https://www.ncbi.nlm.nih.gov/pmc/articles/PMC9943720/ (Accessed July 28, 2023).

Cheng H, Concepcion GT, Feng X, Zhang H, Li H. 2021. Haplotype-resolved de novo assembly using phased assembly graphs with hifiasm. Nat Methods 18: 170–175.

Cheng H, Qu H, McKenzie S, Lawrence KR, Windsor R, Vella M, Park PJ, Li H. 2026. Efficient near-telomere-to-telomere assembly of nanopore simplex reads. Nature 1–8. https://www.nature.com/articles/s41586-026-10105-6 (Accessed February 10, 2026).

Danecek P, Bonfield JK, Liddle J, Marshall J, Ohan V, Pollard MO, Whitwham A, Keane T, McCarthy SA, Davies RM, et al. 2021. Twelve years of SAMtools and BCFtools. Gigascience 10: giab008. 10.1093/gigascience/giab008 (Accessed February 10, 2026).

Del Gobbo GF, Boycott KM. 2025. The additional diagnostic yield of long-read sequencing in undiagnosed rare diseases. Genome Res 35: 559–571. https://pmc.ncbi.nlm.nih.gov/articles/PMC12047273/ (Accessed February 6, 2026).

Garg S, Martin M, Marschall T. 2016. Read-based phasing of related individuals. Bioinformatics 32: i234–i242.

Hofmeister RJ, Ribeiro DM, Rubinacci S, Delaneau O. 2023. Accurate rare variant phasing of whole-genome and whole-exome sequencing data in the UK Biobank. Nat Genet 55: 1243–1249. https://www.nature.com/articles/s41588-023-01415-w (Accessed February 6, 2026).

Kolmogorov M, Billingsley KJ, Mastoras M, Meredith M, Monlong J, Lorig-Roach R, Asri M, Alvarez Jerez P, Malik L, Dewan R, et al. 2023. Scalable Nanopore sequencing of human genomes provides a comprehensive view of haplotype-resolved variation and methylation. Nat Methods 20: 1483–1492. https://www.nature.com/articles/s41592-023-01993-x (Accessed October 10, 2023).

Li H. 2018. Minimap2: pairwise alignment for nucleotide sequences. Bioinformatics 34: 3094–3100.

Li H, Bloom JM, Farjoun Y, Fleharty M, Gauthier L, Neale B, MacArthur D. 2018. A synthetic-diploid benchmark for accurate variant-calling evaluation. Nat Methods 15: 595–597.

Li Q, Keskus AG, Wagner J, Izydorczyk MB, Timp W, Sedlazeck FJ, Klein AP, Zook JM, Kolmogorov M, Schatz MC. 2025. Unraveling the hidden complexity of cancer through long-read sequencing. Genome Res 35: 599–620. https://pmc.ncbi.nlm.nih.gov/articles/PMC12047254/ (Accessed February 10, 2026).

Li W, Freudenberg J. 2014. Mappability and read length. Front Genet 5. https://www.frontiersin.org/journals/genetics/articles/10.3389/fgene.2014.00381/full (Accessed February 6, 2026).

Lin J-H, Chen L-C, Yu S-C, Huang Y-T. 2022. LongPhase: an ultra-fast chromosome-scale phasing algorithm for small and large variants. Bioinformatics 38: 1816–1822. 10.1093/bioinformatics/btac058 (Accessed April 17, 2023).

Martin M, Patterson M, Garg S, Fischer SO, Pisanti N, Klau GW, Schöenhuth A, Marschall T. 2016. WhatsHap: fast and accurate read-based phasing. 085050. https://www.biorxiv.org/content/10.1101/085050v2 (Accessed February 6, 2026).

Miller DE, Lee L, Galey M, Kandhaya-Pillai R, Tischkowitz M, Amalnath D, Vithlani A, Yokote K, Kato H, Maezawa Y, et al. 2022. Targeted long-read sequencing identifies missing pathogenic variants in unsolved Werner syndrome cases. J Med Genet 59: 1087–1094.

Miller DE, Sulovari A, Wang T, Loucks H, Hoekzema K, Munson KM, Lewis AP, Fuerte EPA, Paschal CR, Walsh T, et al. 2021. Targeted long-read sequencing identifies missing disease-causing variation. The American Journal of Human Genetics 108: 1436–1449. https://www.sciencedirect.com/science/article/pii/S0002929721002305 (Accessed April 6, 2023).

Mosbruger TL, Dinou A, Duke JL, Ferriola D, Li Y, Hayeck TJ, Monos DS. 2025. Haplotypic resolution of the challenging genomic regions of MHC and KIR using a combination of targeted sequencing and a novel assembly pipeline. Nucleic Acids Res 53: gkaf441. 10.1093/nar/gkaf441 (Accessed February 10, 2026).

Nakamura H, Doi H, Mitsuhashi S, Miyatake S, Katoh K, Frith MC, Asano T, Kudo Y, Ikeda T, Kubota S, et al. 2020. Long-read sequencing identifies the pathogenic nucleotide repeat expansion in RFC1 in a Japanese case of CANVAS. J Hum Genet 65: 475–480. https://www.nature.com/articles/s10038-020-0733-y (Accessed August 3, 2023).

Nurk S, Koren S, Rhie A, Rautiainen M, Bzikadze AV, Mikheenko A, Vollger MR, Altemose N, Uralsky L, Gershman A, et al. 2022. The complete sequence of a human genome. Science 376: 44–53. https://pmc.ncbi.nlm.nih.gov/articles/PMC9186530/ (Accessed February 9, 2026).

Nyaga DM, Zaied RE, Silander OK, Black MA, O’Sullivan JM. 2025. Beyond single references: pangenome graphs and the future of genomic medicine. Front Genet 16: 1679660. https://pmc.ncbi.nlm.nih.gov/articles/PMC12492951/ (Accessed April 21, 2026).

Pedregosa F, Varoquaux G, Gramfort A, Michel V, Thirion B, Grisel O, Blondel M, Prettenhofer P, Weiss R, Dubourg V, et al. 2011. Scikit-learn: Machine Learning in Python. Journal of Machine Learning Research 12: 2825–2830. http://jmlr.org/papers/v12/pedregosa11a.html (Accessed April 7, 2026).

Samarasinghe SR, Gaedigk A, Swen JJ, Guchelaar H-J, Nagaraj SH. 2026. Long-Read Sequencing Enhances Pharmacogenomic Profiling by Resolving Complex Haplotypes, Novel Star Alleles, and Structural Variants. Clin Pharmacol Ther 119: 536–545.

Schmitz D, Ameur A, Johansson Å. 2025. T2T-CHM13 improves read mapping and detection of clinically relevant genetic variation in the Swedish population. Genome Res 35: 2377–2388. https://pmc.ncbi.nlm.nih.gov/articles/PMC12581843/ (Accessed February 10, 2026).

Shafin K, Pesout T, Chang P-C, Nattestad M, Kolesnikov A, Goel S, Baid G, Kolmogorov M, Eizenga JM, Miga KH, et al. 2021. Haplotype-aware variant calling with PEPPER-Margin-DeepVariant enables high accuracy in nanopore long-reads. Nat Methods 18: 1322–1332. https://www.nature.com/articles/s41592-021-01299-w (Accessed April 10, 2023).

Snyder MW, Adey A, Kitzman JO, Shendure J. 2015. Haplotype-resolved genome sequencing: experimental methods and applications. Nat Rev Genet 16: 344–358. https://www.nature.com/articles/nrg3903 (Accessed February 9, 2026).

Tewhey R, Bansal V, Torkamani A, Topol EJ, Schork NJ. 2011. The importance of phase information for human genomics. Nat Rev Genet 12: 215–223. https://www.nature.com/articles/nrg2950 (Accessed February 6, 2026).

van der Lee M, Rowell WJ, Menafra R, Guchelaar H-J, Swen JJ, Anvar SY. 2022. Application of long-read sequencing to elucidate complex pharmacogenomic regions: a proof of principle. Pharmacogenomics J 22: 75–81. https://www.nature.com/articles/s41397-021-00259-z (Accessed August 8, 2023).

Wang Q, Pierce-Hoffman E, Cummings BB, Alföldi J, Francioli LC, Gauthier LD, Hill AJ, O’Donnell-Luria AH, Karczewski KJ, MacArthur DG. 2020. Landscape of multi-nucleotide variants in 125,748 human exomes and 15,708 genomes. Nat Commun 11: 2539. https://pmc.ncbi.nlm.nih.gov/articles/PMC7253413/ (Accessed March 26, 2026).

Zheng Z, Li S, Su J, Leung AW-S, Lam T-W, Luo R. 2022. Symphonizing pileup and full-alignment for deep learning-based long-read variant calling. Nat Comput Sci 2: 797–803. https://www.nature.com/articles/s43588-022-00387-x (Accessed April 10, 2023).

