## Supplemental Figures for "Determinants of haplotype phasing accuracy in long-read human genome sequencing"

**Table of Contents**

**Figure S1.** **Read length distributions for ONT R10 Simplex and PacBio HiFi from HG002 and HG005.**

**Figure S2.** Comparison of phasing error rates between alignment-based and assembly-based phasing methods.

**Figure S3.** IGV screenshots of *ARHGEF18* in HG002 and HG005 showing phaseset assignment across technologies, samples and reference genomes.

**Figure S4**. Variant **pair counts and phasing error rates for ARHGEF18 in HG002 and HG005.**

**Figure S5.** IGV screenshots of *RYR2* in HG002 showing phaseset assignment across technologies, reference genomes and phasing methods.

**Figure S6.** Comparison of phasing error rate in *HYDIN* in HG002 across different phasing methods.

**Figure S7.** IGV screenshots of *HYDIN* in HG002 showing phaseset assignment across technologies, reference genomes and phasing methods.

**Figure S8.** Phasing error rate versus number of variant pairs (log scale) for individual OMIM genes in HG002.

**Figure S9**. Phasing error rate versus number of variant pairs (log scale) for individual OMIM genes in HG002.

**Figure S10.** Gene-level phasing error rates for HG002 aligned to GRCh38 and T2T, comparing alignment-based and assembly-based phasing for ONT and PacBio.

**Figure S11.** Gene-level phasing error rates for HG005 aligned to GRCh38 and T2T, comparing alignment-based and assembly-based phasing for ONT and PacBio.

**Figure S12. Random forest feature importance for predicting phasing errors across technologies and reference genomes.**

**Figure S1. Read length distributions for ONT R10 Simplex and PacBio HiFi from HG002 (left) and HG005 (right).** Histograms show the distribution of primary aligned read lengths (≥100 bp) for ONT R10 Simplex (blue) and PacBio HiFi (orange) reads aligned to GRCh38 (top row) and T2T-CHM13v2 (bottom row). Dashed vertical lines indicate the read N50 for each dataset.

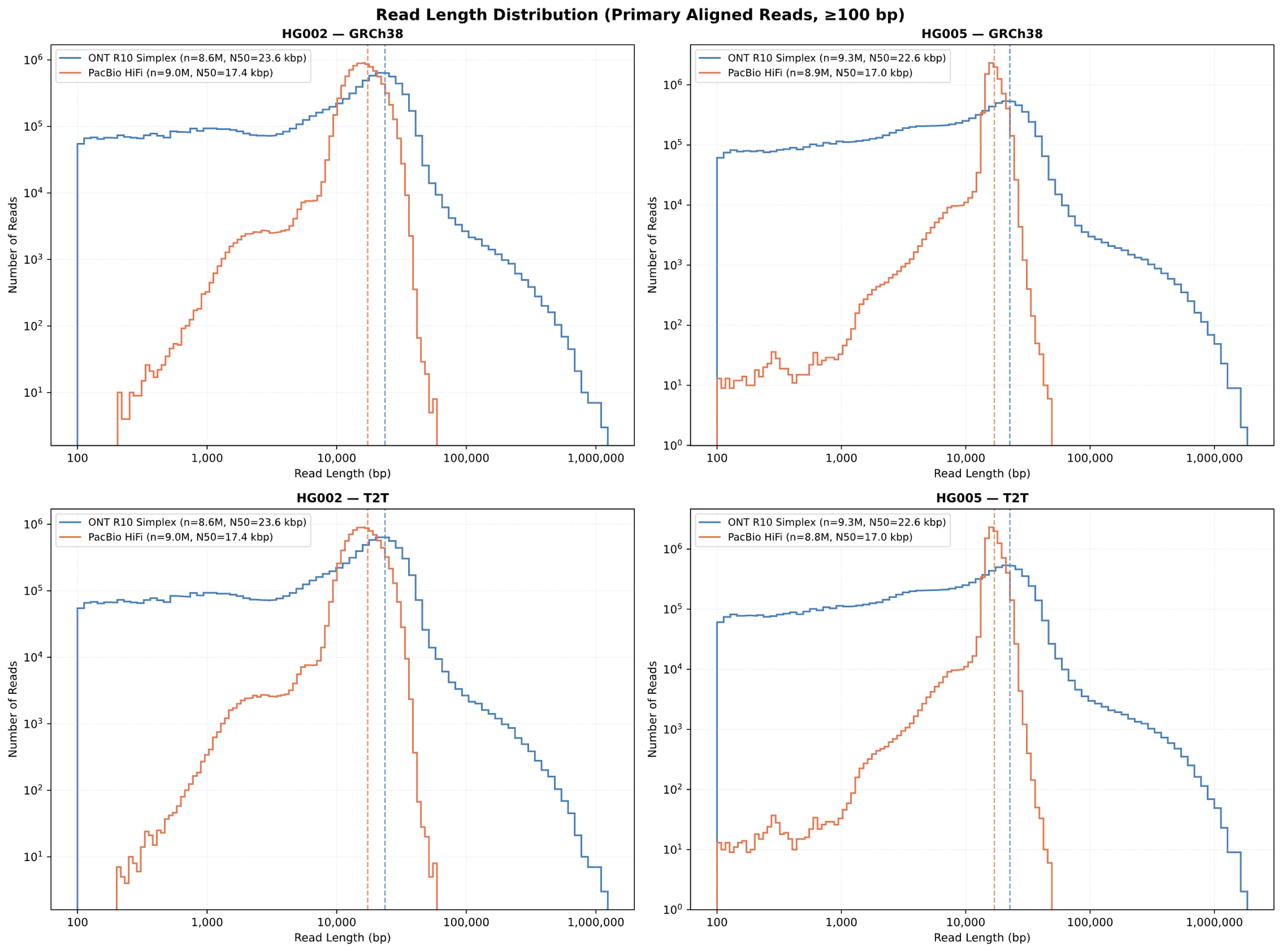

**Figure S2. Comparison of phasing error rates between alignment-based and assembly-based phasing methods.** **(A,B)** Number of high-quality phased variant pairs in HG002 **(A)** and HG005 **(B)** by distance between variants for ONT and PacBio using alignment-based phasing and haplotype-resolved de novo assembly, stratified by reference genome and GIAB-defined genomic regions. **(C,D)** Phasing error rates by distance between variants in HG002 **(C)** and HG005 **(D)** for ONT and PacBio using alignment-based phasing and haplotype-resolved de novo assembly, stratified by reference genome and GIAB-defined genomic regions.

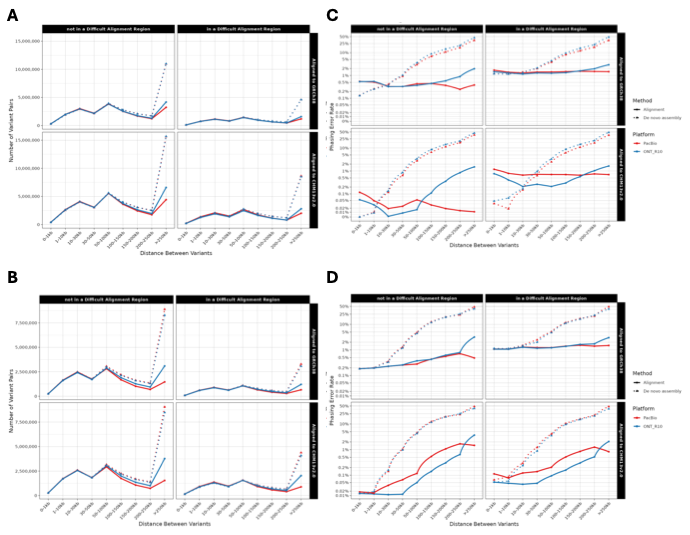

**Figure S3.** IGV screenshots of *ARHGEF18* in HG002 and HG005 showing phaseset assignment across technologies, samples and reference genomes. Top panel shows GRCh38 alignments; bottom panel shows T2T-CHM13v2.0 alignments. Reads are colored by haplotype assignment (PS tag).

GRCh38

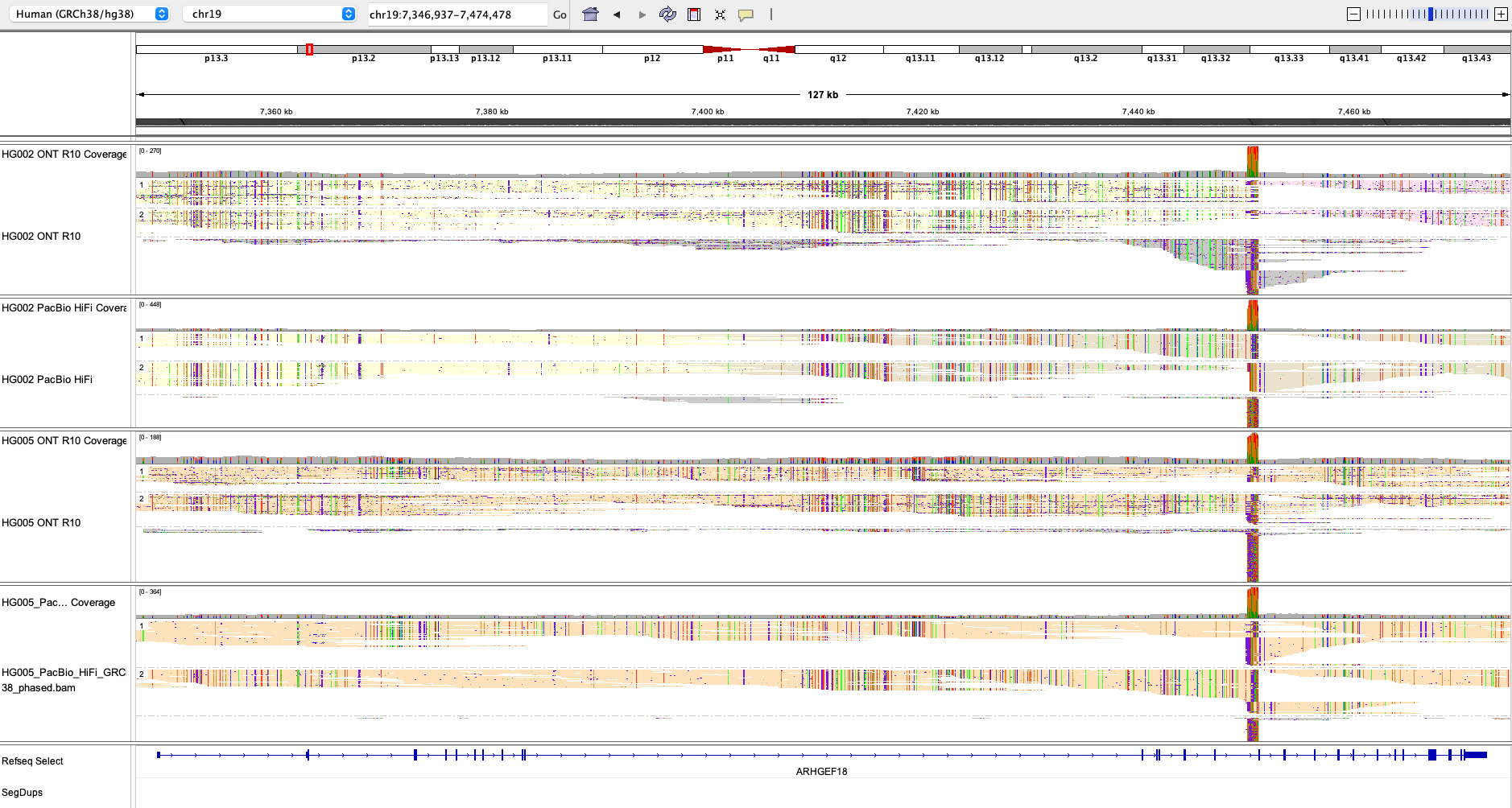

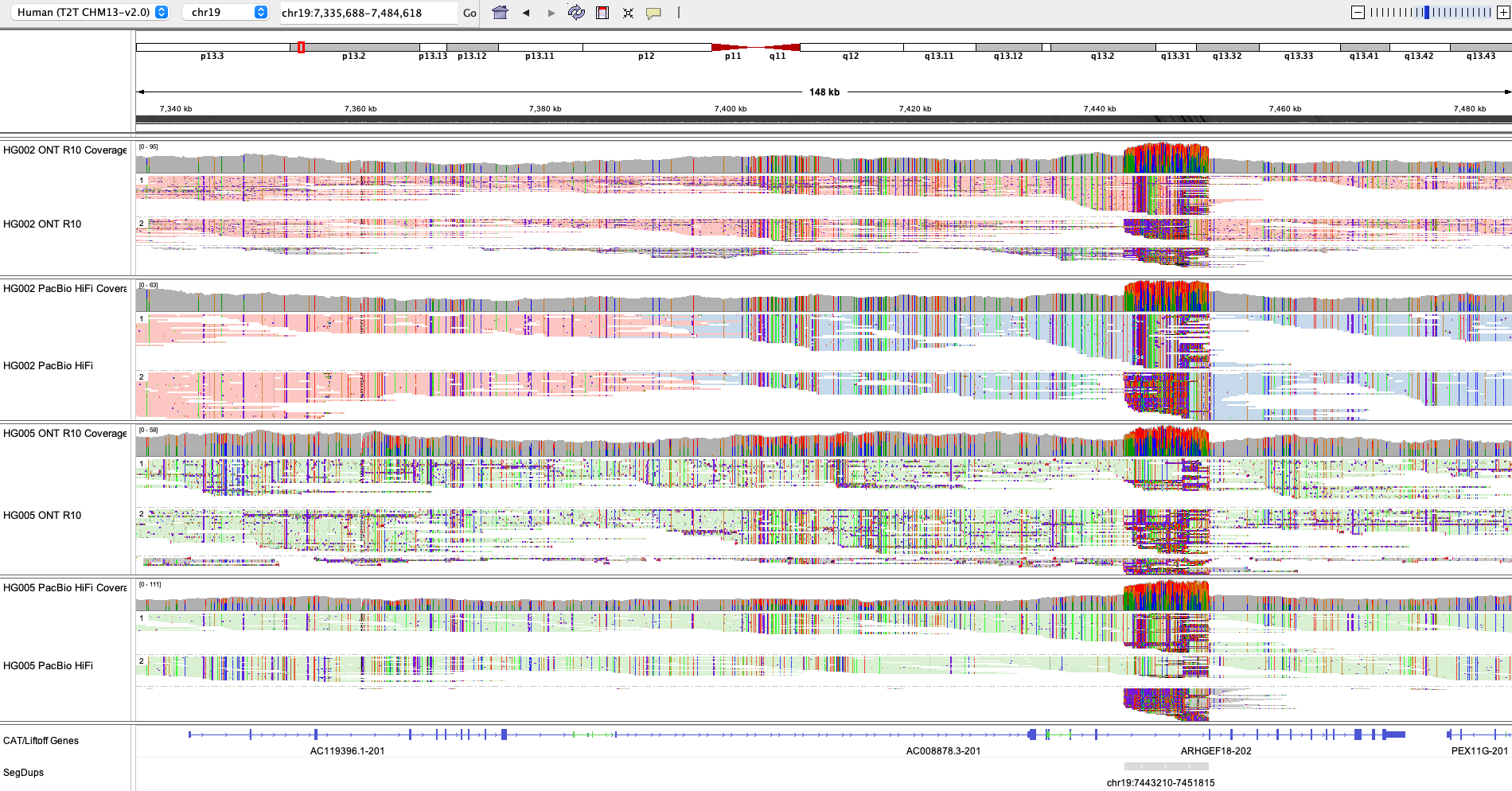
T2T

**Figure S4. Variant pair counts and phasing error rates for ARHGEF18 in HG002 and HG005.** Variant pair counts (top) and phasing error rates (bottom) are shown across distance bins for ONT R10 and PacBio HiFi aligned to GRCh38 and T2T CHM13v2.0, stratified by difficult alignment regions.

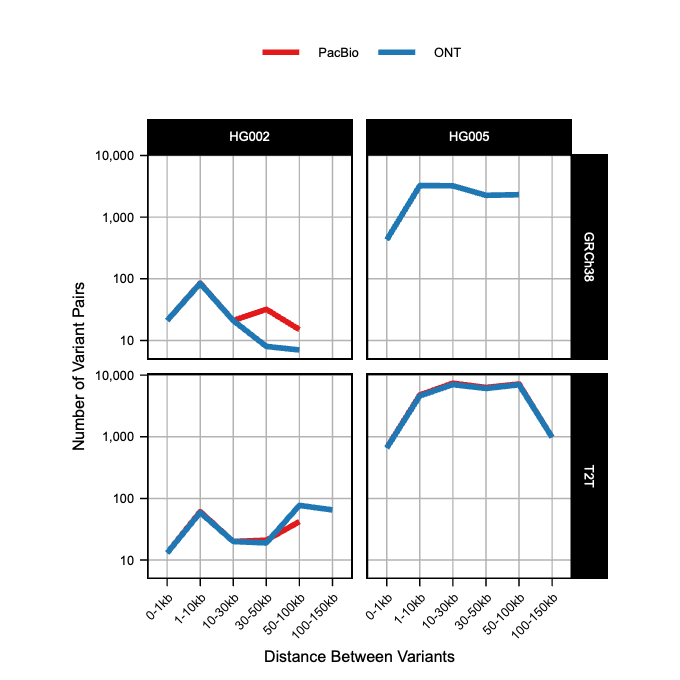

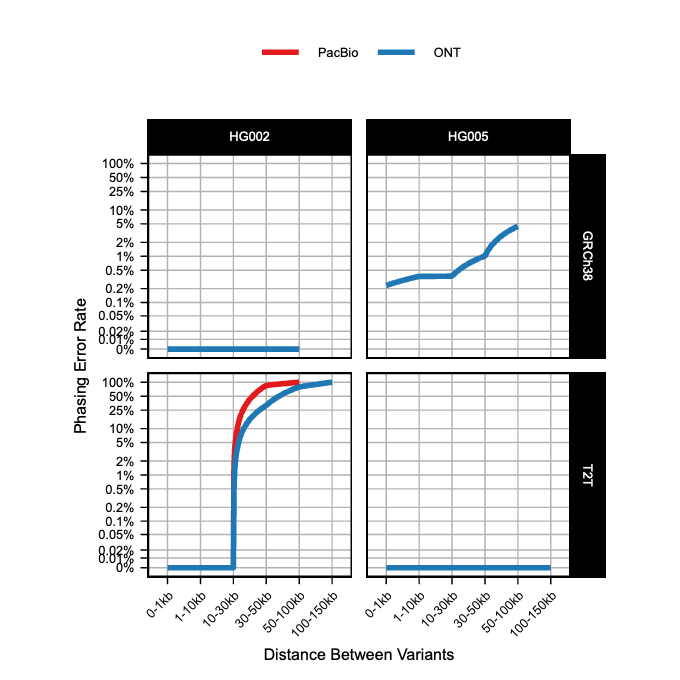

**Figure S5. IGV screenshots of *RYR2* in HG002 showing phaseset assignment across technologies, reference genomes and phasing methods.** Each screenshot displays multiple tracks representing different combinations of sequencing technology (ONT, PacBio HiFi) and phasing method (Longphase, PMDV, WhatsHap). Reads are colored by haplotype assignment.

GRCh38

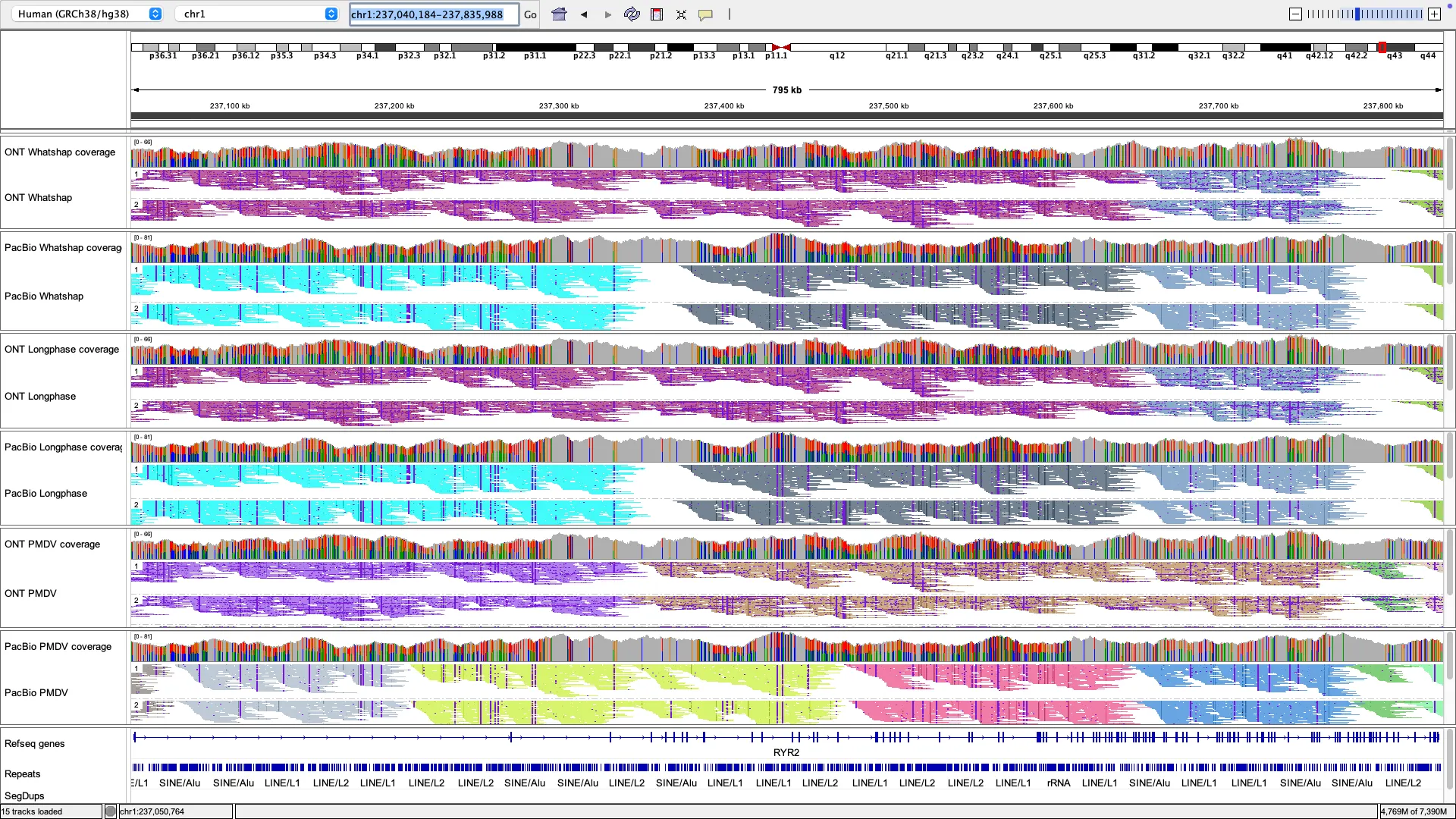

T2T

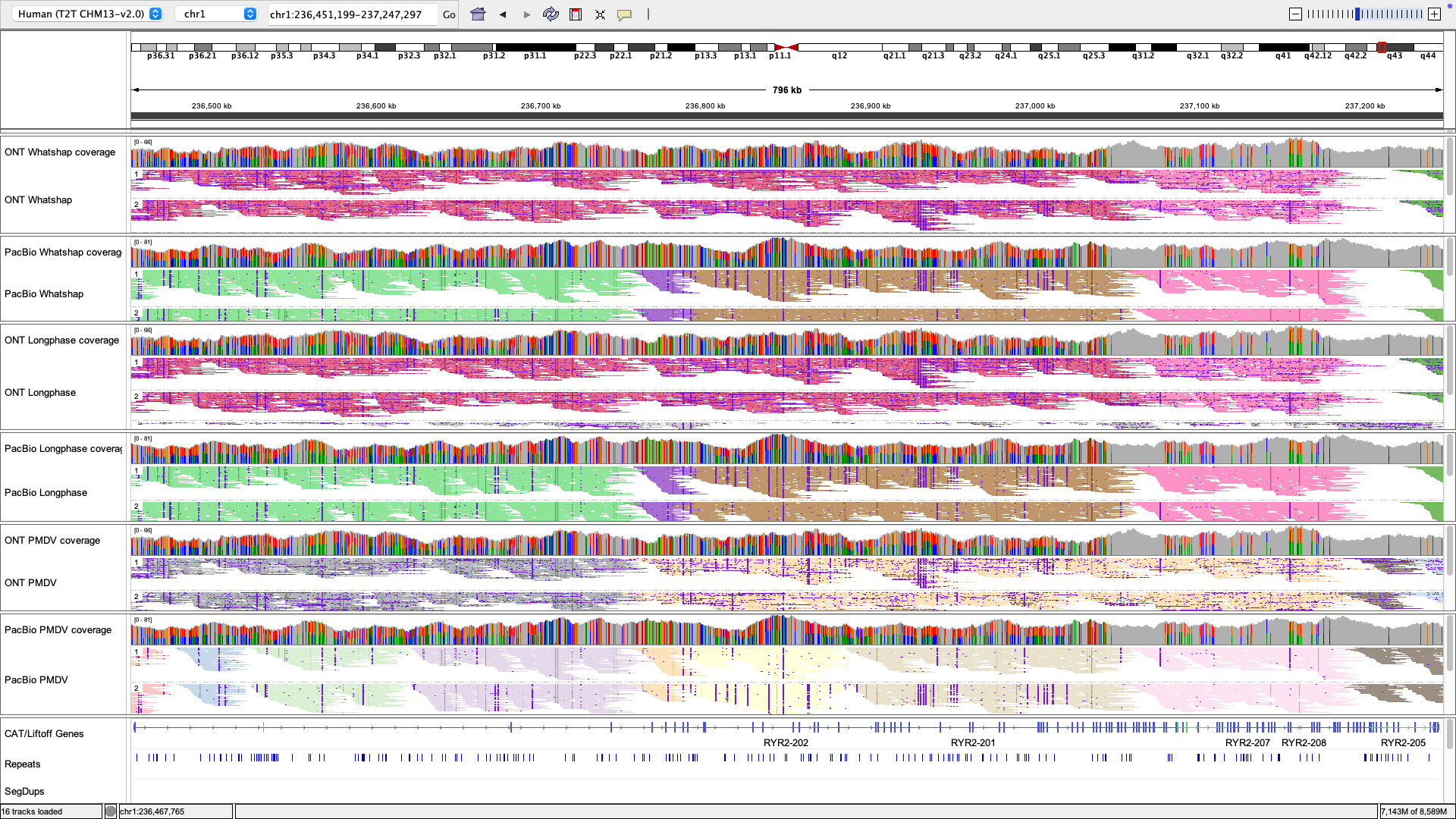

**Figure S6. Comparison of phasing error rate in *HYDIN* in HG002 across different phasing methods.** Top panel shows the total number of variant pairs phased in *HYDIN* for each method and technology combination. Bottom panel shows phasing error rates for *HYDIN* using three phasing methods (Longphase, PMDV, and WhatsHap) for ONT (left) and PacBio HiFi (right).

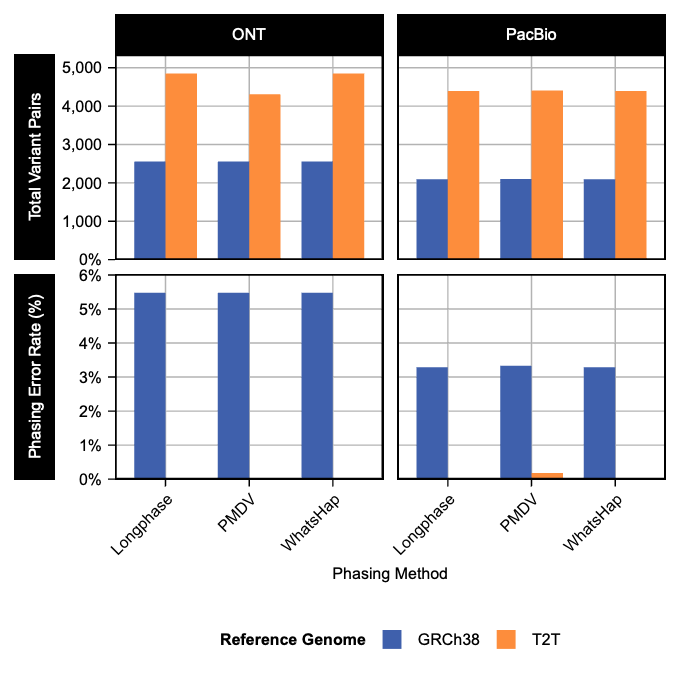

**Figure S7. IGV screenshots of *HYDIN* in HG002 showing phaseset assignment across technologies, reference genomes and phasing methods**. Top panel shows GRCh38 alignments; bottom panel shows T2T alignments. Each screenshot displays multiple tracks representing different combinations of sequencing technology (ONT, PacBio HiFi) and phasing method (Longphase, PMDV, WhatsHap). Reads are colored by haplotype assignment.

GRCh38

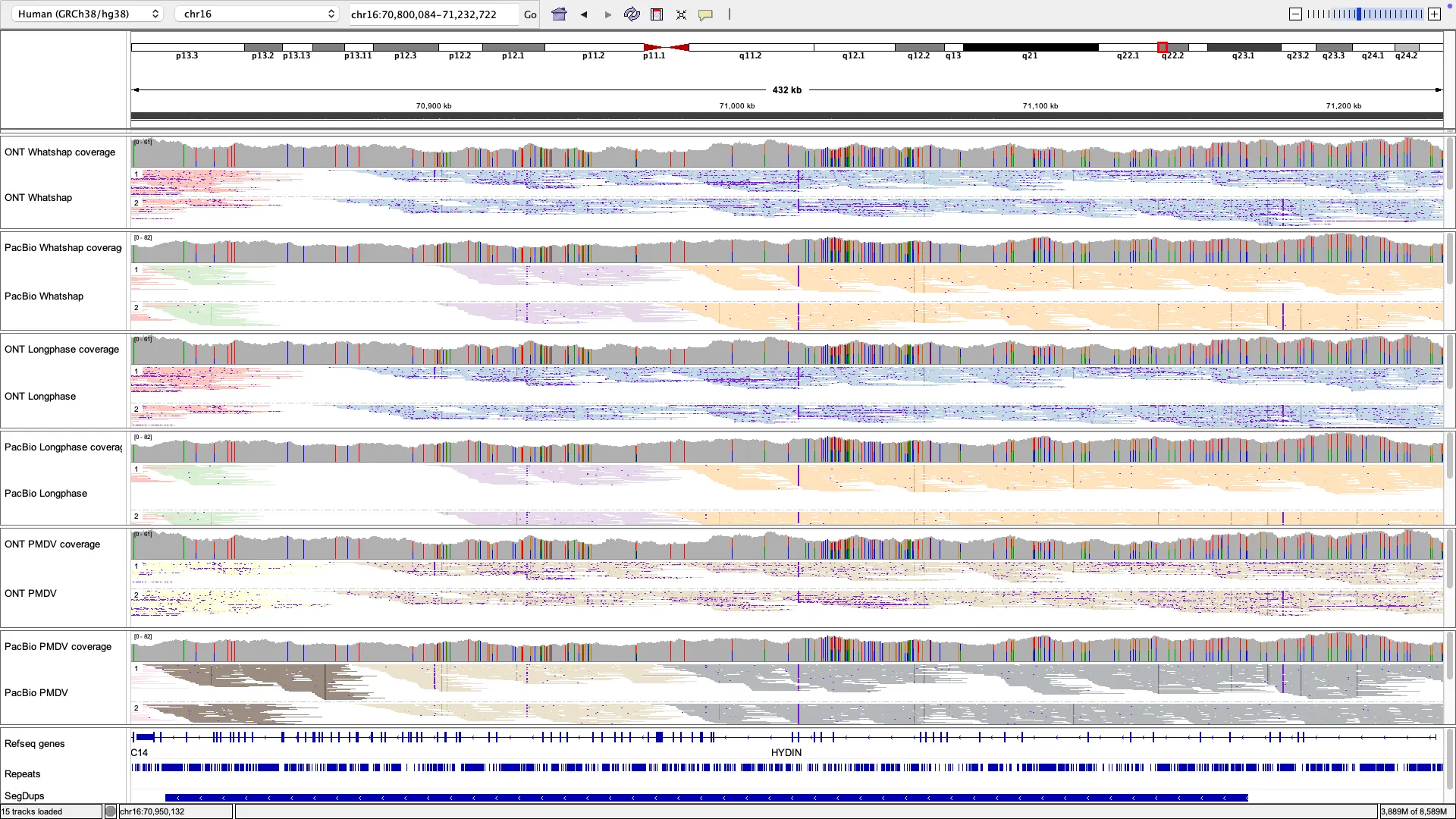

T2T

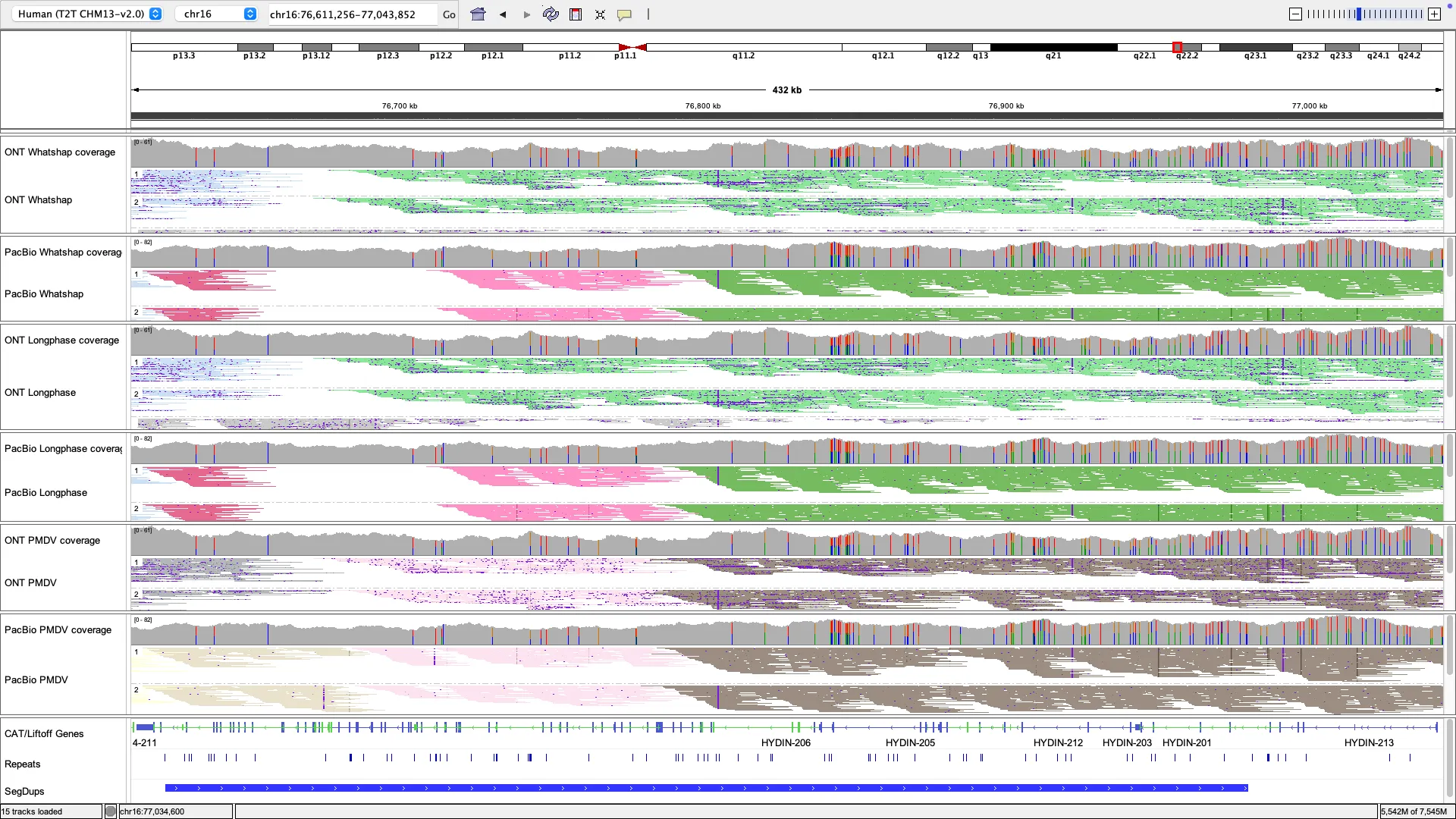

**Figure S8. Phasing error rate versus number of variant pairs (log scale) for individual OMIM genes in HG002 using ONT R10 data.** Top panels show alignment-based phasing results for GRCh38 (left) and T2T-CHM13v2.0 (right). Bottom panels show haplotype-resolved *de novo* assembly (hifiasm) results for GRCh38 (left) and T2T-CHM13v2.0 (right). Select genes with elevated error rates are labeled. The horizontal dashed line indicates 50% phasing error rate.

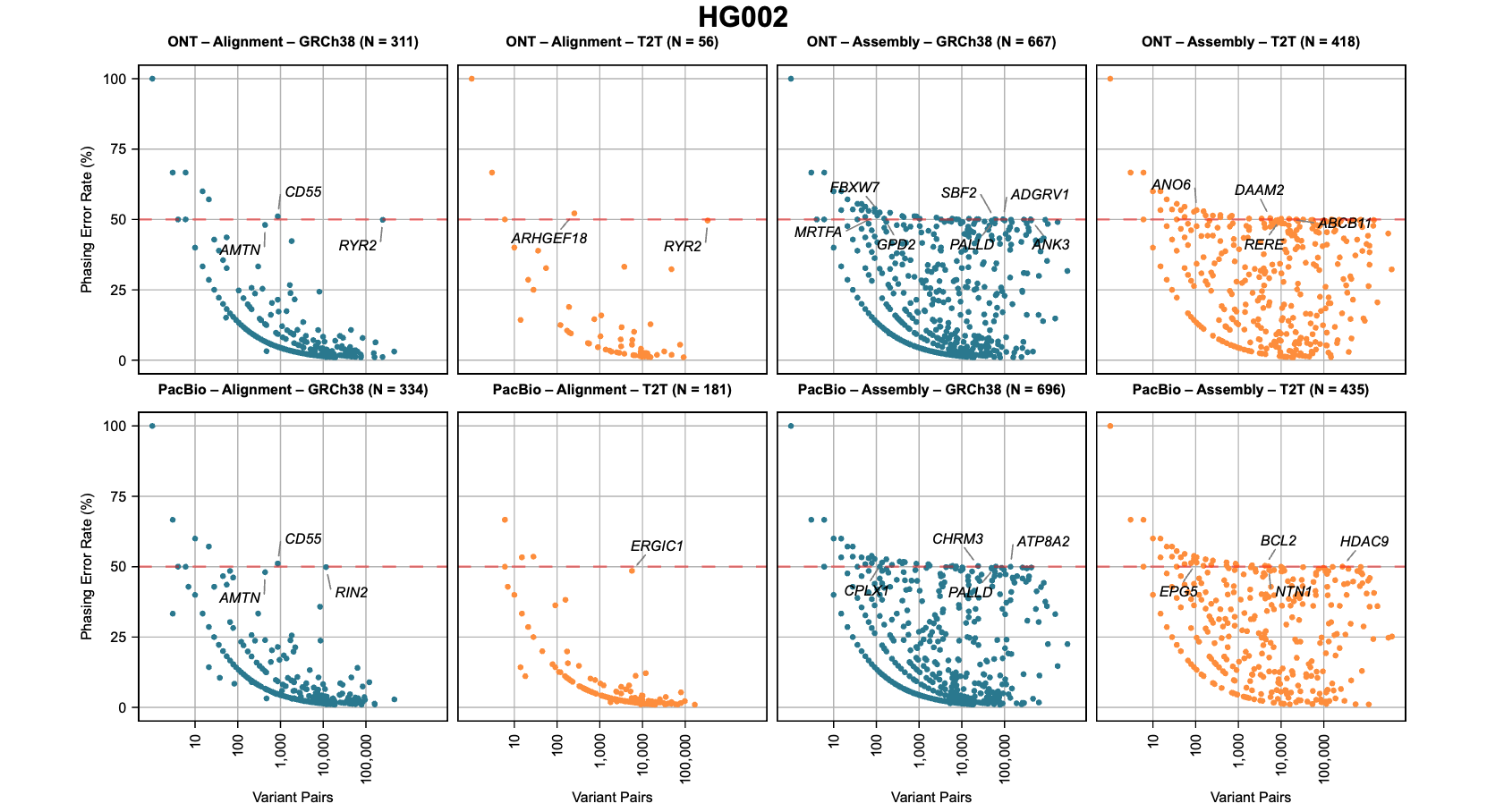

**Figure S9. Phasing error rate versus number of variant pairs (log scale) for individual OMIM genes in HG005 using PacBio HiFi data.** Top panels show alignment-based phasing results for GRCh38 (left) and T2T (right). Bottom panels show haplotype-resolved *de novo* assembly (hifiasm) results for GRCh38 (left) and T2T (right). Select genes with elevated error rates are labeled. The horizontal dashed line indicates 50% phasing error rate.

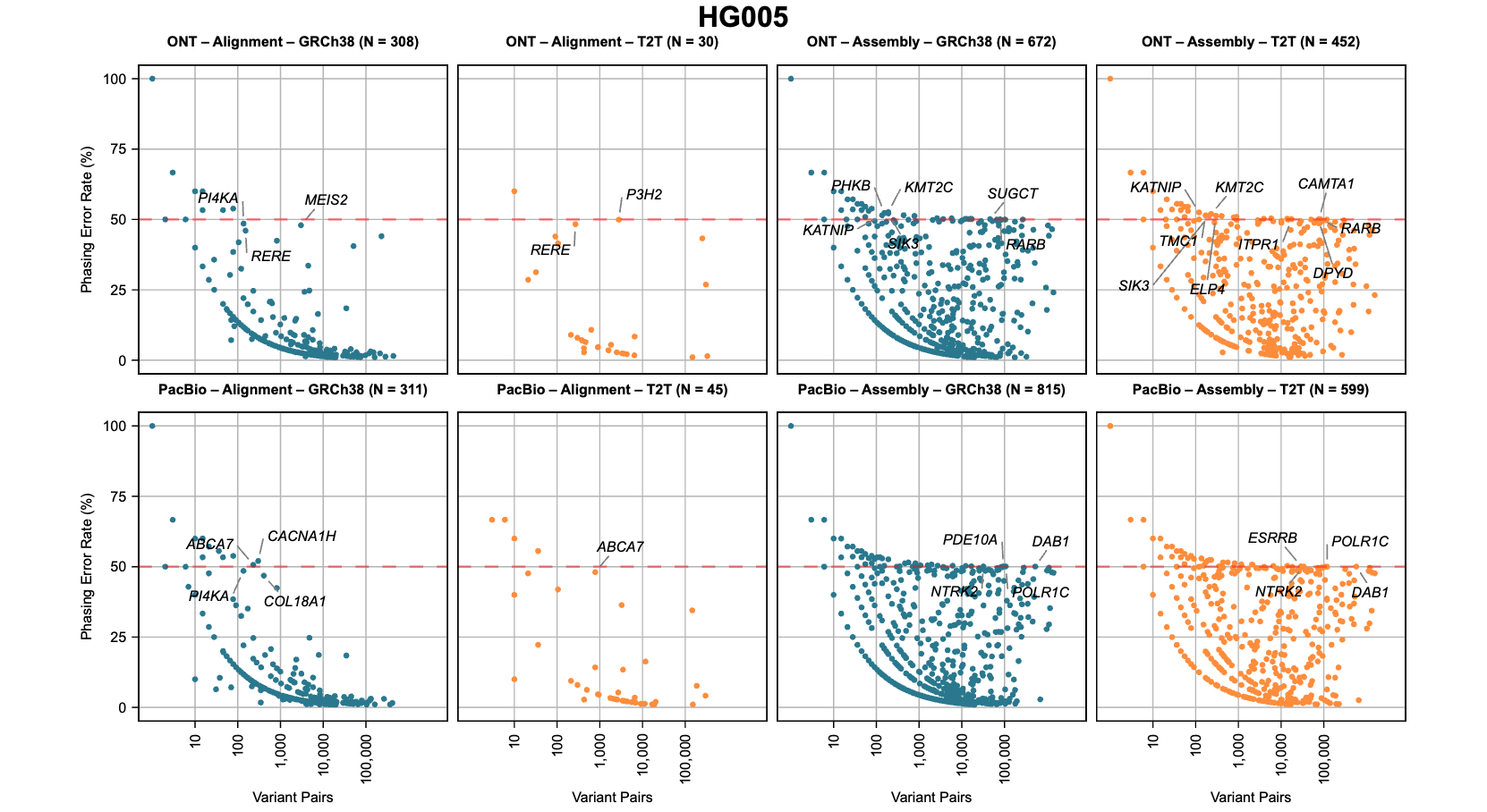

**Figure S10. Gene-level phasing error rates for HG002 aligned to GRCh38 and T2T, comparing alignment-based and assembly-based phasing for ONT and PacBio.** Each panel shows genes with a phasing error rate ≥50% and >100 variant pairs in at least one method (alignment or assembly) for that technology-reference genome combination. Genes shown in bold are common across all four panels. Genes are ordered by descending maximum error rate within each technology panel.

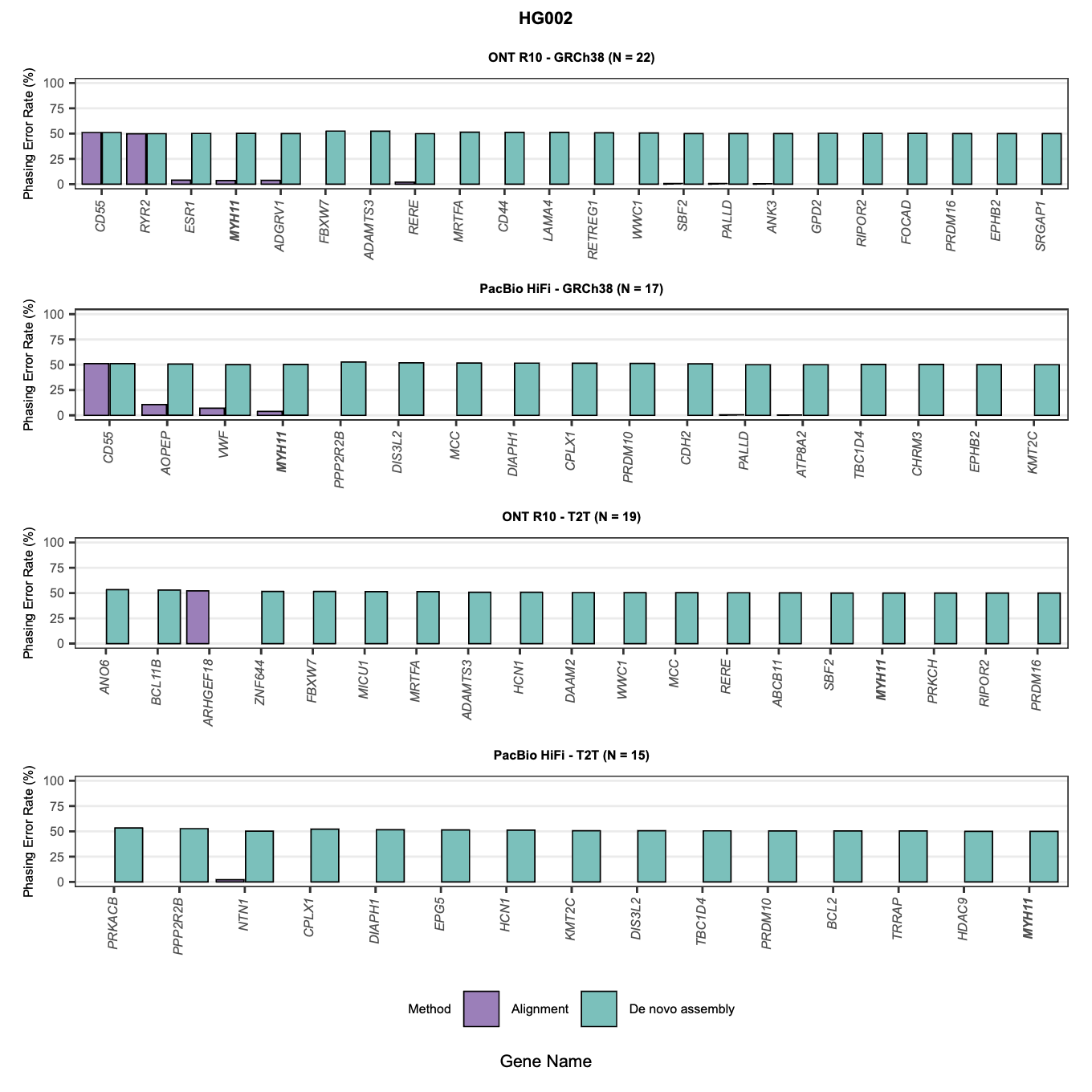

**Figure S11. Gene-level phasing error rates for HG005 aligned to GRCh38 and T2T, comparing alignment-based and assembly-based phasing for ONT and PacBio.** Each panel shows genes with a phasing error rate ≥50% and >100 variant pairs in at least one method (alignment or assembly) for that technology-reference genome combination. Genes shown in bold are common across all four panels. Genes are ordered by descending maximum error rate within each technology panel**.**
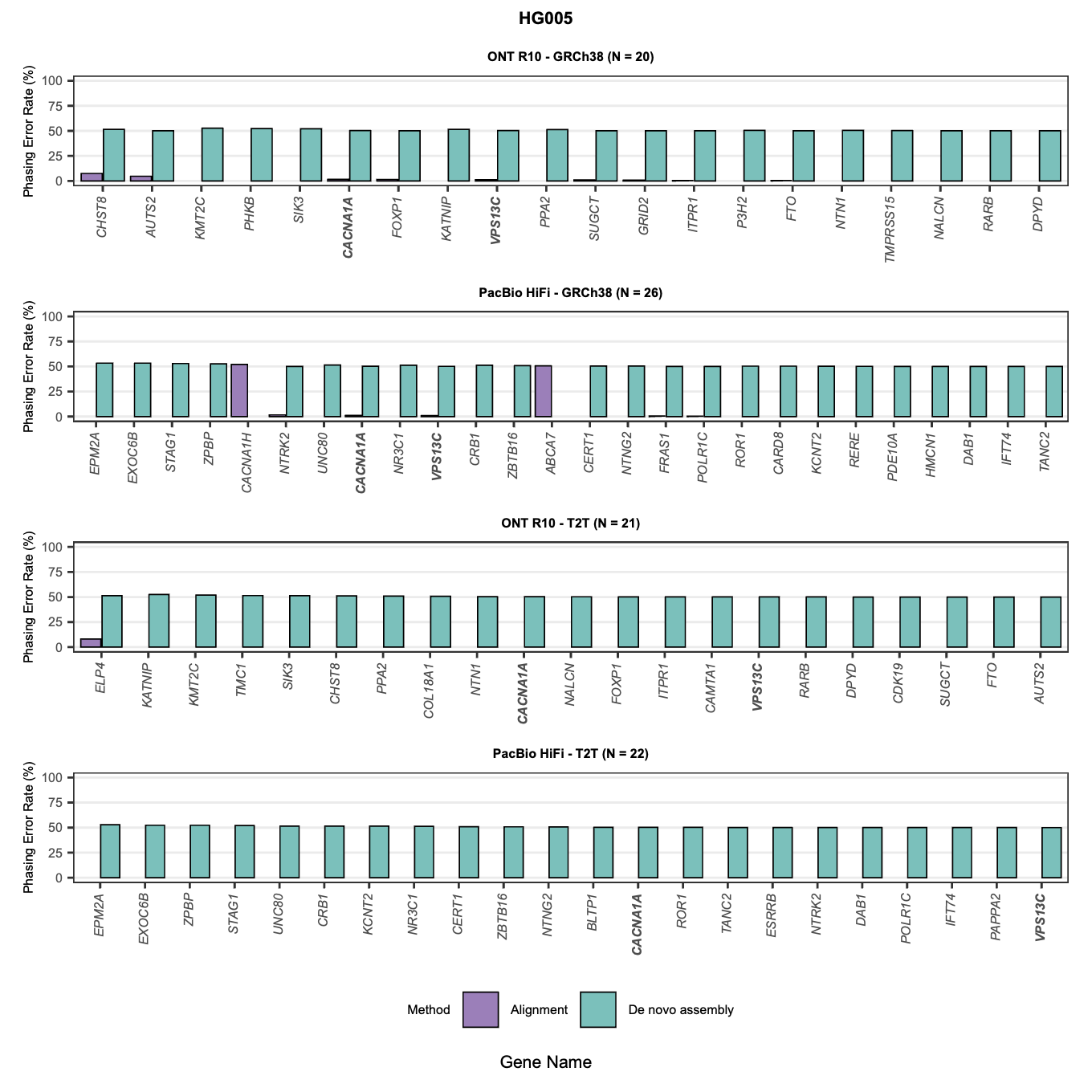

**Figure S12. Random forest feature importance for predicting phasing errors across technologies and reference genomes.** Feature importance scores from random forest classifiers trained on a subset of HG002 variant pairs. Models were trained separately for each technology (ONT R10, PacBio HiFi) and reference genome (GRCh38, T2T) combination. Features include: difficult-to-align region (overlap with segmental duplications or low mappability regions), maximum gap between variants (largest gap in base pairs within the variant pair span), phase set size (size of the phase block truncated to gene boundaries), gap-to-read ratio (maximum gap divided by longest aligned read in block), maximum read in phase set (longest aligned read length in the phase block), and variant density (number of variants per kilobase between the pair). Higher importance values indicate features that are more informative for distinguishing correct from incorrect phasing in the training dataset.

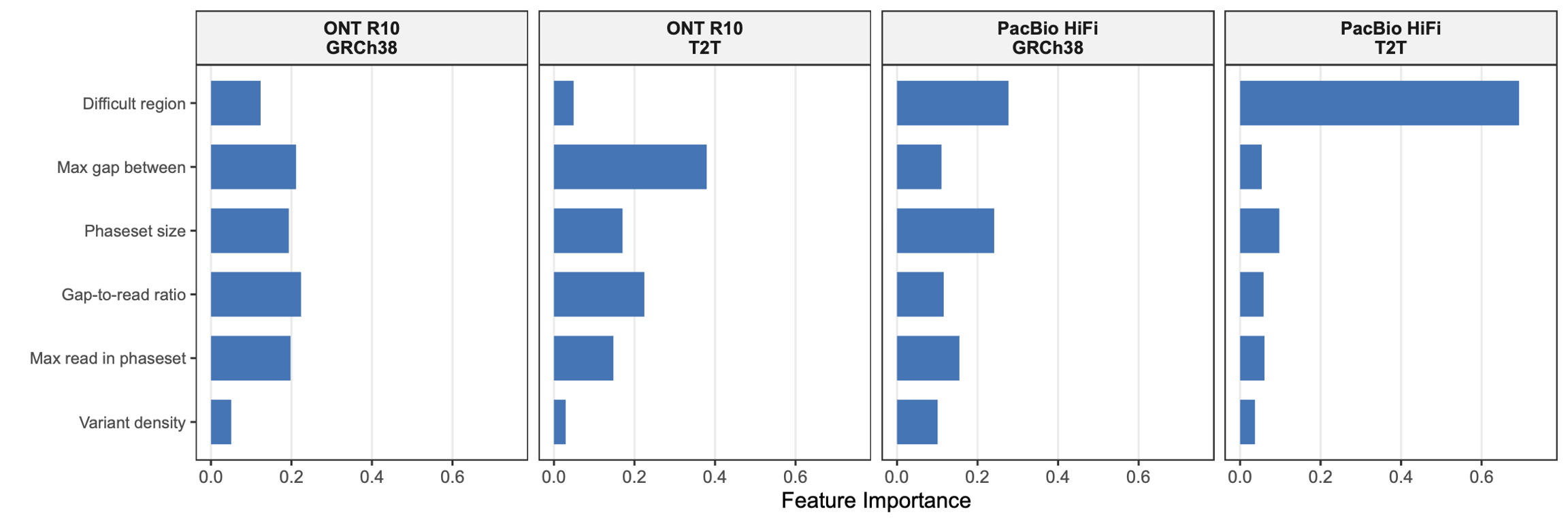
